# Volumetric imaging and single-cell RNAseq atlases identify cellular mechanisms of human dental pulp response during tooth decay progression

**DOI:** 10.1101/2025.05.11.653296

**Authors:** Hoang-Thai Ha, Sofya Kosmynina, Amandine Verocq, Keremsah Ozen, Ines Tekia, Hugo Bussy, Dima Sabbah, Chloe Goemans, Valerie Vandenbempt, Esteban Gurzov, Sumeet Pal Singh, Nicolas Baeyens

## Abstract

Dental pulp responses to dental decay, the most prevalent chronic disease worldwide, involve remodeling processes similar to those observed in other human pathological conditions. By integrating volumetric imaging and single-cell analysis across different disease stages in human samples, we uncovered the natural history of dental pulp responses to decay. At early stages, we observed an arterialization of the capillary networks and progressive outward remodeling of the larger vessels. Neurogenesis of nerve endings and the reprogramming of perivascular progenitor cells into fibroblasts are also observed, initiating the physiological reparative response of the stroma. Pathological angiogenesis and nerve regression combined with dental pulp fibrosis at later stages of tooth decay determine irreversible pulpitis. These results provide a basis for understanding dental tissue response to injury, driving a paradigm shift in patient management.

**Graphical abstract:** 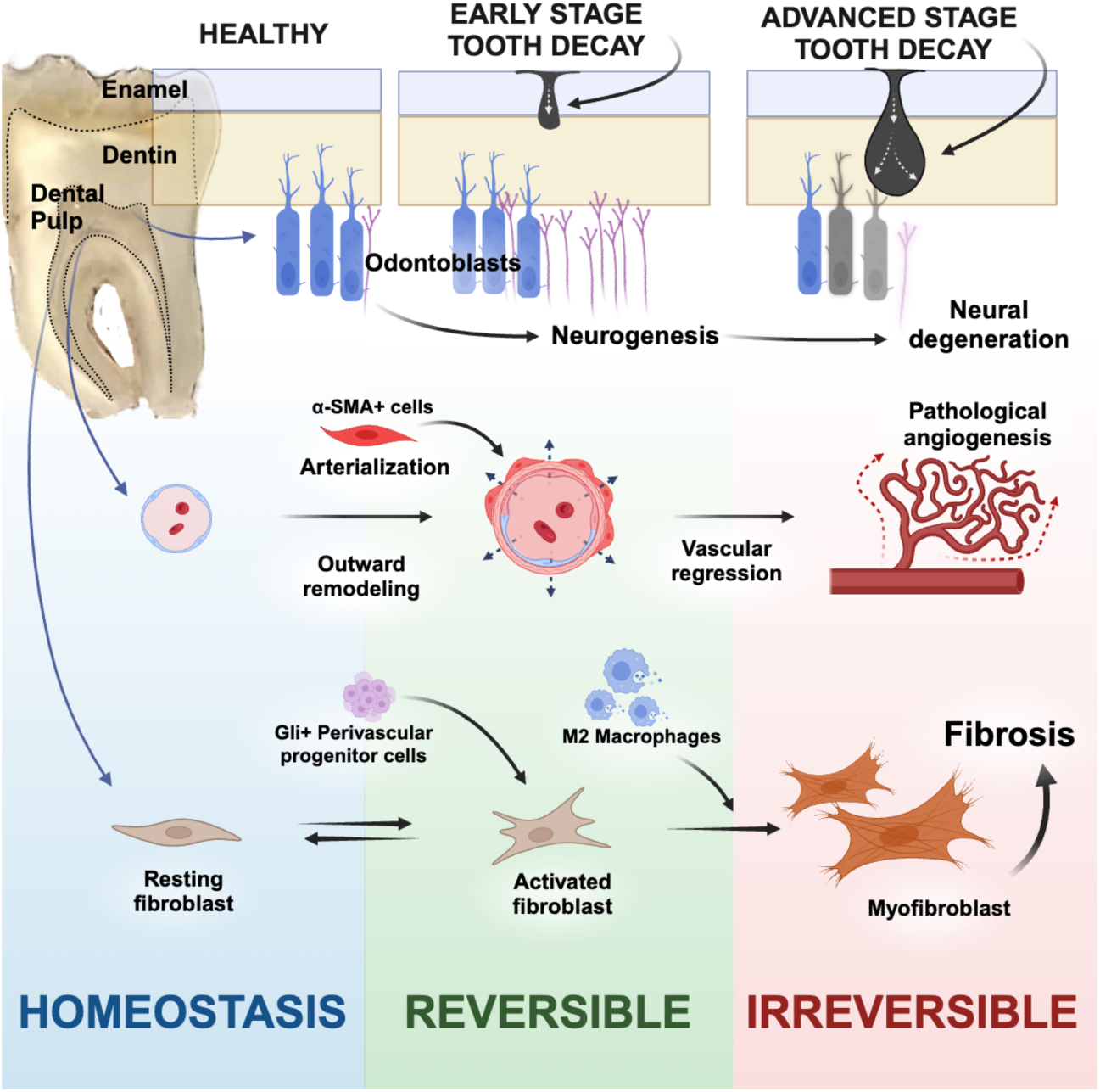

## Introduction

Dental decay is a slowly progressing infectious disease, the most common chronic disease worldwide (*1*). It causes enamel demineralization, followed by cavitation and subsequent progression of decay into the dentin. This progression can be clinically and histologically classified, as done by the SiSta classification (*2*), based on the progression of the decay inside the hard tissue. The decay of the hard tissues releases metabolic byproducts into the dentinal tubules, which can diffuse to the dentin-pulp interface. Pulpal cells sense invading pathogens and trigger a cascade of inflammatory responses called pulpitis (*3*). Current clinical practices focus on removing infected hard tissues with limited consideration for the pulp’s regenerative potential. The cellular and molecular mechanisms involved in pulp response remain unclear, mainly how these processes vary with disease progression and severity and what drives the progression from healthy, reversible pulpitis into pathological, irreversible pulpitis. While a pilot study has demonstrated increased expression of pro-inflammatory, anti-inflammatory, and mineralization-related genes in cases of deep decay (*4*), a deeper understanding of these mechanisms during the progression of the disease is necessary. Precise insights could drive a paradigm shift in dental treatment, transitioning from hard tissue management and pulp removal by endodontic approaches to strategies targeting pulp preservation and regeneration.

The spatial and functional characteristics of the cellular systems in the dental pulp, along with its heterogeneity and response to dental decay, remain largely unexplored due to the unusual properties of the tooth. The dental pulp is a specialized soft connective tissue located at the tooth’s core within a rigid, opaque, non-expandable chamber. It is encased by hard, mineralized tissues: an outer enamel layer and an inner dentin layer comprising dentinal tubules, allowing for communication between the outer layers of the hard tissue and the pulp. The pulp is highly innervated and vascularized, with its vascular system playing a vital role in tooth homeostasis by facilitating cell nutrition, oxygenation, and the removal of metabolic waste (*5*). Meanwhile, the neural system regulates blood flow, tooth sensitivity, and pain perception (*6*). Previous research using resin casts (*7*) and Indian ink (*8*) in animal models has revealed the basics of the pulp vasculature’s three-dimensional architecture. More recently, a study imaged the neurovascular systems of the human dental pulp extracted from hard tissues after tissue clearing (*5*). However, these studies were either limited to animal models that do not naturally develop tooth decay as observed in humans or involved pulp extracted from its mineralized environment, often with limited imaging resolution. Visualization at high resolution of the complete architecture of the neurovascular system in human teeth, with a preserved dentin-pulp interface, is lacking. The human tooth is unique because it contains fully functional soft tissue that is accessible and easily harvested, for example, during wisdom teeth extraction, enabling procedures such as omics investigations and therapeutic trials.

In this work, we developed a comprehensive atlas of the natural history of tooth decay by combining volumetric imaging of cleared human tooth samples with a preserved dentin-pulp interface and single-cell analysis of human dental pulps in both healthy and diseased states. This approach allowed us to provide detailed insights into the remodeling processes as the pathology progresses. We identified the potential for reversibility in early disease stages, followed by irreversible remodeling as the disease progresses to advanced stages if no dental care is undertaken. The various remodeling processes observed are also present in other systemic diseases, highlighting the potential of the human tooth as a unique and promising model for studying human systemic conditions.

## Results

### Volumetric Imaging Unveils the Global Architecture of the Neurovascular Systems in the Healthy Human Dental Pulp: Innervation and Perfusion are Aimed at the Dentin-Pulp Interface

To investigate the spatial characteristics of the human dental pulp, samples from healthy (n=40) and diseased (n=74) teeth were collected, processed into thick sections of >1mm, preserving the dentin-pulp interface, sections were cleared with a modified iDISCO protocol (*9*), immunolabeled, and imaged across the whole surface of the pulp at a high-resolution with a confocal microscope, by fusing several z-stacks together (Figure 1A). The dental pulp exhibits a dense, well-organized vasculature (Figure 1B, S1). High-resolution 3D volumetric images of blood vessels (Figure 1C) and maximum intensity projections (MIP) from z-stacks were generated. We identified five distinct blood vessel types in the dental pulp based on size (Figure S1B), location (Figure S1A), endothelial cell (EC) alignment, and mural cell (smooth muscle cells [SMCs] and pericytes) coverage. Feeding arterioles from the tooth roots merge centrally, branching into vertical arterioles that supply a dense capillary plexus beneath the dentin-pulp interface. Blood drains through vertical venules into larger collecting venules, exiting via the roots. Endothelial cells (ECs), marked by vascular endothelial (VE) cadherin-specific antibodies targeting an endothelial-specific adhesion molecule found at EC junctions (*10*), respond to blood flow forces known as blood flow-induced, fluid, or wall shear stress (*11, 12*), which influence their number, shape and alignment (*13*). Arteries experience higher shear stress (10–70 dynes/cm²), promoting EC elongation, while veins, with lower stress (1–6 dynes/cm²), exhibit a cobblestone-like morphology (Figure 1D-1E) (*12, 13*). Structurally, arteries have thick tunica media layers with multiple smooth muscle cell (SMC) layers, while venules have thinner, less resistant walls. Capillaries, consisting of a single EC layer with no tunica media, form a dense plexus at the dentin-pulp interface, highlighting their vital role in pulp vascularization (Figure 1F). Mural cells were identified using antibodies targeting α-smooth muscle actin (*14*). The presence of lymphatic vessels within the dental pulp has been debated for many years (*15*). In our study, immunostaining with lymphatic-specific antibodies such as LYVE1, PROX1, and D2-40 (Podoplanin) were used but did not reveal the presence of a lymphatic network within healthy dental pulp (Figure S4A-S4C). Our study provides a comprehensive and detailed volumetric characterization of the dental pulp vasculature.

**Figure 1.**
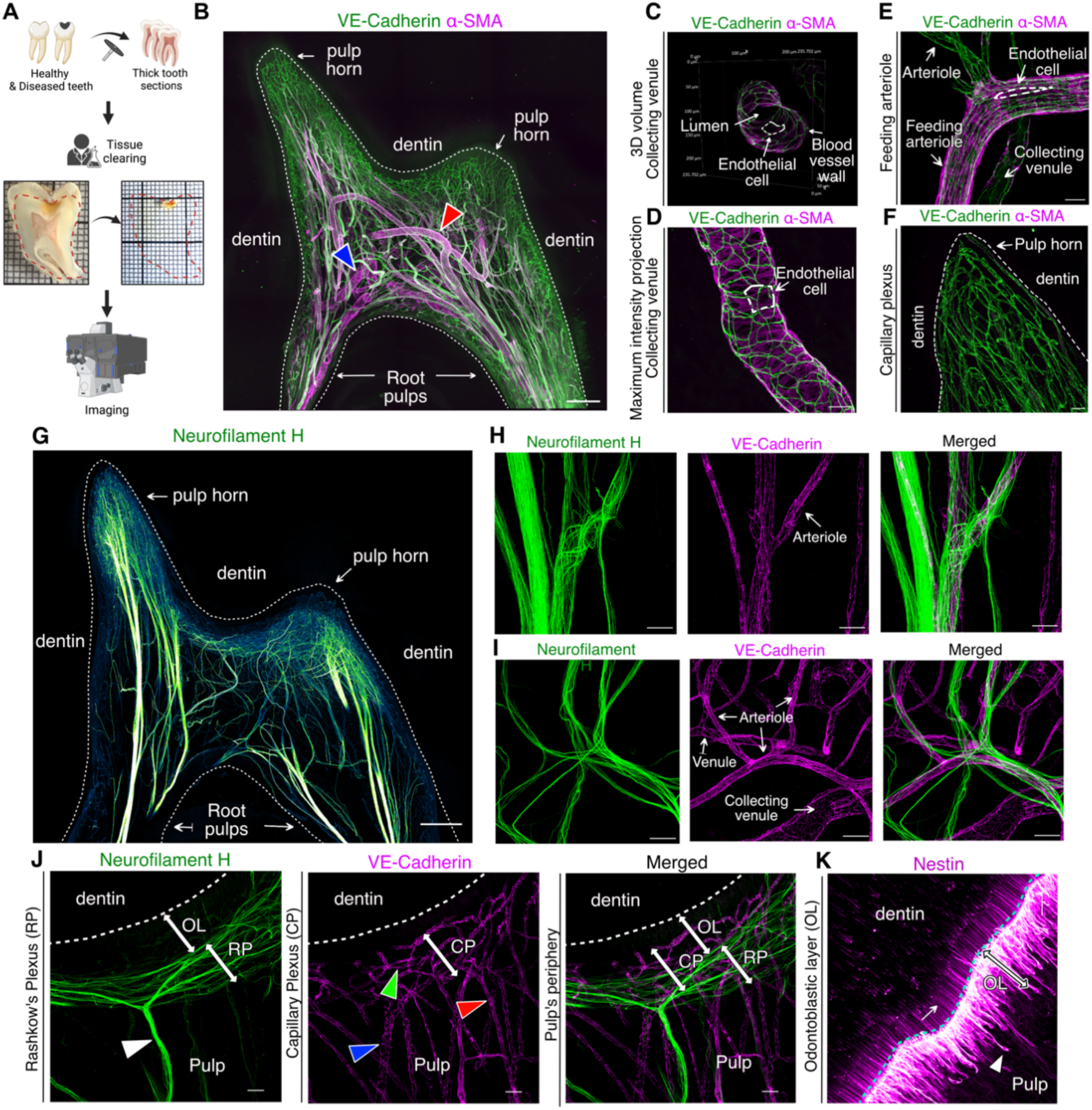
**Global Architecture of the Neurovascular Systems in the Healthy Human Dental Pulp** (A) Schematic of the imaging experimental workflow. (B) Maximum intensity projection (MIP) of a 343μm z-stack showing the vasculature of a healthy mature dental pulp. (red arrow) a feeding arteriole; (blue arrow) a collecting venule. (C) 3D projection of a 98µm-stack showing a collecting venule. (D) MIP of a 100µm z-stack showing a collecting venule. (E) MIP of a 79µm z-stack showing a feeding arteriole. (F) MIP of a 302µm z-stack showing capillaries. (white dashed line) an endothelial cell. (G) MIP of a 252 μm z-stack showing the neural network of a healthy mature dental pulp. (H-J) (H)MIP of a 118µm z-stack and (I) a 143µm z-stack showing the close neurovascular interactions, (J) a 143µm-stack showing the neurovascular interaction at the periphery of the pulp. (white arrowhead) a nerve trunk; (Blue arrowhead) a venule; (red arrowhead) an arteriole; (Green arrowhead) a capillary. (K) MIP of a 91µm z-stack showing the odontoblastic layer. Tissue clearing using a modified iDISCO protocol and staining for the vasculature (B-F) with VE-Cadherin (green) and α-SMA (purple), for the neural network (G-J) with Neurofilament-H (green) and VE-Cadherin (purple), and for the odontoblastic cells (K) with Nestin (purple). Point scanning resonant confocal microscope. Large images for (B,G). Plan Apo 10x λS OFN25 DIC N1 optic for (B, D, G, I, J). Plan apo Lambda 25XC Sil. optic for (C, E, F, H, K). Scale bars: 500µm for (B,G), 100µm for (I), 50µm for (F, H, J), 25µm for (D, E, K).

Nerve fibers were immunolabeled with Neurofilament H (NF-H)-specific antibodies (*16*), revealing a dense network. Large nerve trunks enter the root apex, traverse the roots, and branch extensively in the pulp chamber before reaching the periphery (Figure 1G). Neurovascular interactions, well-described in peripheral tissues (*17*) and dental pulp (*18*), involve unmyelinated C-fibers (regulating blood flow, dull pain, and running along blood vessels) and myelinated A-fibers (mediating tooth sensitivity, sharp pain, and running independently) (Figures 1H, 1I and Video S1) (*18*). At the periphery near the dentin-pulp interface, nerve trunks form a dense Raschkow’s plexus (RP) under the odontoblastic layer (OL), closely interacting with the capillary plexus (CP) (Figure 1J). Occasionally, some nerve fibers extend beyond the RP, crossing the OL to penetrate the dentin via dentinal tubules (Figures 1J, S2A and S2B).

Odontoblasts, specialized cells at the pulp periphery, reside in the odontoblastic layer (OL) with their cell bodies adjacent to the dentin. Their processes extend into the dentin via tubules to varying depths. As the outermost layer of the dental pulp, odontoblasts lie above the Raschkow’s plexus (Figures 1K, S2A-C). Odontoblasts play crucial roles: they produce secondary and tertiary dentin in response to stimuli or injury and facilitate pulpal sensory functions through molecular crosstalk with nerve fibers. The mechanisms underlying these interactions remain unclear. The dense capillary plexus right under the OL ensures metabolic support, which is essential for these functions.

### Cellular Heterogeneity of the Human Dental Pulp

To comprehensively map the cellular heterogeneity of human dental pulp, we constructed a transcriptomic atlas using a split-pool combinatorial barcoding single-cell RNA sequencing protocol. We analyzed healthy (n=4), early-stage diseased (n=4), and advanced-stage diseased (n=4) human pulp samples from molar teeth extracted for clinical reasons (Table S1). Tissue samples were extracted, dissociated into single-cell suspensions, fixed using Parse fixation kits, and stored at −80°C. All samples were processed using the Parse Evercode® WT v2 kit, sequenced, and analyzed using Trailmaker® (Figure 2A). After filtering, integrating (Harmony method), and clustering (Louvain method), we analyzed 53,407 cells. We identified 12 main clusters representing five systems: fibroblast and stromal cells (51% of the cell population), endothelial cells (ECs) (30% of the cell population), immune and blood cells (8% of the cell population), mural cells (5% of the cell population), and glial cells (5% of the cell population) (Figures 2B, 2C, S3A and S3B). A signature of *CXCL14, LUM, C1S, DCN, COL1A1* and *COL1A2* characterized the fibroblast and stromal cells cluster (*19, 20*). ECs were identified with the following canonical markers: *CDH5*, *VWF*, *EMCN*, *ENG,* and *FLT4* (*21, 22*), while immune and blood cells were identified with the expression of *FTL*, *CD68*, *MS4A7*, *MAF*, *TYROBP*, *CD74*, *PTPRC,* and *SKAP1* (*23*). Mural cells specifically expressed high levels of *ACTA2*, *TAGLN*, *BGN*, *NOTCH3*, *RGS5*, *PDGFRB* and *MCAM* (*20*). Finally, glial cells expressed *MPZ*, *MBP*, *PMP22*, *DMD*, *TRPM3*, *XKR4*, *ADGRB3,* and *GRIK2* (Figures 2D, 2E and S3C-G) (*24*).

**Figure 2.**
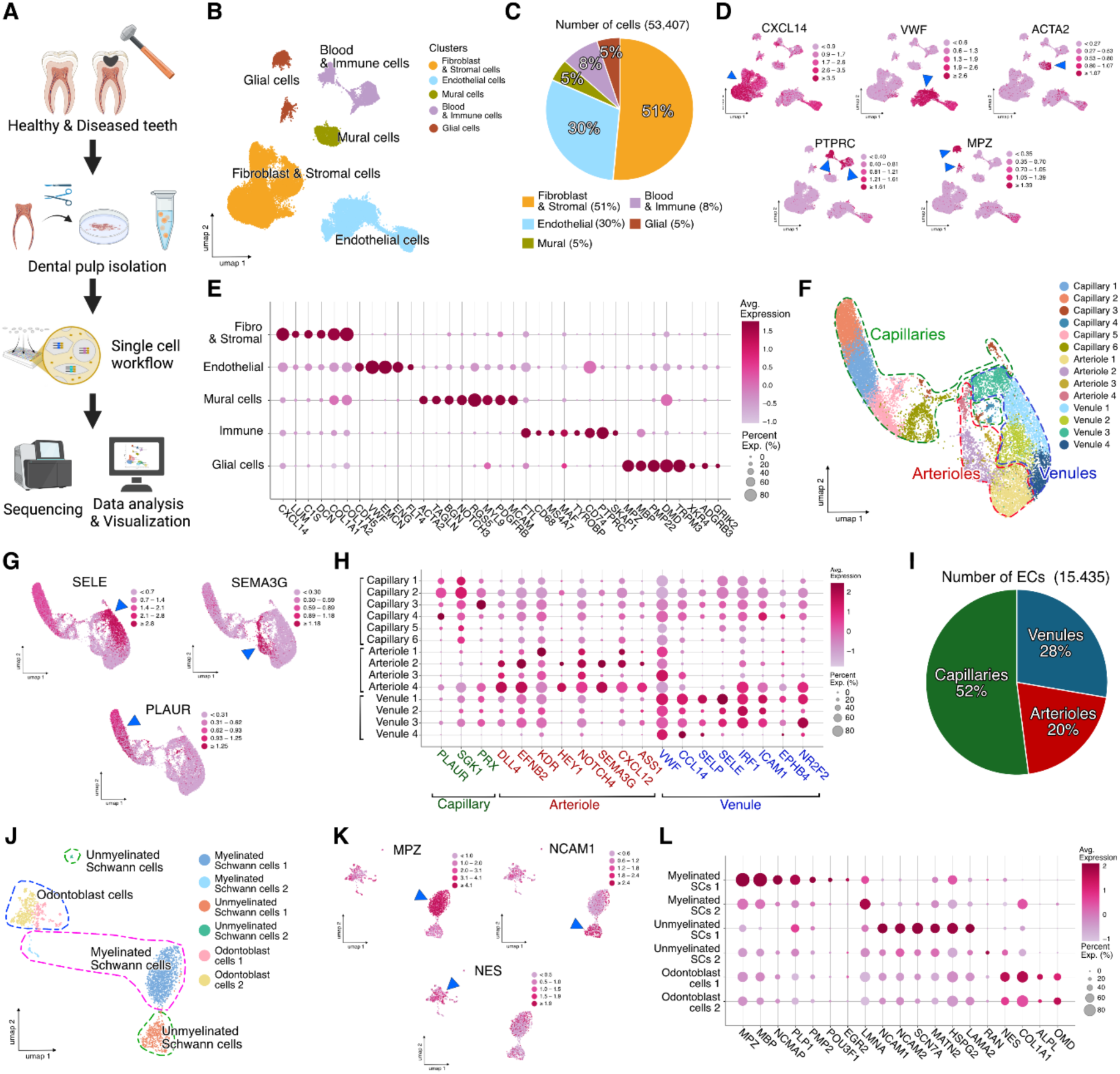
**Cellular Heterogeneity of the Human Dental Pulp** (A) Schematic of the single-cell experimental workflow. (B) Uniform manifold approximation and projection (UMAP) representation of 53.407 filtered and batch corrected healthy and diseased dental pulp cells across 4 healthy pulps, 4 early stage and 4 advanced stage diseased pulps. (C) Proportions of each cell population. (D) Relative expression of a well-established cell type marker for each cell population projected on UMAP plots. *CXCL14* for fibroblast and stromal cells; *VWF* for ECs; *ACTA2* for mural cells; *PTPRC* for immune cells; *MPZ* for Glial cells. (E) Dot plot showing 5-8 well-established cell type markers allowing to identify each cluster. (F) UMAP representation of 15,435 filtered and batch corrected healthy and diseased dental pulp endothelial cells across 4 healthy pulps, 4 early stage and 4 advanced stage diseased pulps. (G) Relative expression of a well-established cell type marker for each EC population projected on UMAP plots. *SELE* for venules; *SEMA3G* for arterioles; *PLAUR* for capillaries. (H) Dot plot showing 3-8 well-established cell type markers allowing to identify each endothelial cells sub-clusters. (I) Graph showing the proportion of each different vascular structures. (J) UMAP representation of 2,440 filtered, and batch corrected healthy and diseased dental pulp glial cells across 4 healthy pulps, 4 early stage and 4 advanced stage diseased pulps. (K) Relative expression of a well-established cell type marker for each glial cells population projected on UMAP plots. *MPZ* for myelinated SCs; *NCAM1* for unmyelinated SCs; *NES* for odontoblast cells. (L) Dot plot showing 4-8 well-established cell type markers allowing to identify each glial cell sub-clusters.

The endothelial cell population was further analyzed. We identified 15.435 endothelial cells divided into fourteen subclusters: six subclusters of capillary endothelial cells (52% of the endothelial cell population) characterized by a higher expression of *PLAUR*, *SGK1* and *PRX* (*21, 25*); four subclusters of arteriolar endothelial cells (20% of the ECs population) characterized with the following markers: *DLL4*, *EFNB2*, *KDR*, *HEY1*, *NOTCH4*, *SEMAG3G*, *CXCL12,* and *ASS1* (*22, 25*); and four subclusters of venous endothelial cells (28% of the ECs population) expressing *VWF*, *CCL14*, *SELP*, *SELE*, *IRF1*, *ICAM1*, *EPHB4* and *NR2F2* (Figures 2F-2I) (*22*). Our single-cell analyses did not reveal any distinct lymphatic subcluster within the endothelial cells, and no significant expression of canonical lymphatic markers such as *LYVE1, PROX1,* or *PDPN* was detected (*26*). However, within the venous subclusters, the venule 1 subcluster exhibited co-expression of *PROX1, ICAM1,* and *VCAM1*, suggesting the presence of functionally specialized reactive post-capillary venules (REVs) (Figure S4D and S4E) (*27*). Our data highlight the crucial role of the vascular system in the dental pulp, particularly emphasizing the predominance of capillaries over other vessel types. This confirms that the dental pulp has a higher average capillary density compared to most other tissues (*28*). The abundance of capillaries suggests that, like skeletal muscles, the dental pulp is a metabolically active tissue.

The glial cells population was further analyzed, identifying 2.440 cells divided into six subclusters: two subclusters of myelinated Schwann cells (SCs) expressing *MPZ*, *MBP*, *NCMAP*, *PLP1*, *POU3F1*, *EGR2* and *LMNA* (*29, 30*); two subclusters of non-myelinated SCs characterized by the expression of *NCAM1*, *NCAM2*, *SCN7A*, *MATN2*, *HSPG2*, *LAMA2* and *RAN* (*29, 30*); and two subclusters of odontoblastic cells expressing *NES*, *COL1A1*, *ALPL* and *OMD* (Figures 2J, 2K and 2L) (*31–33*)*. NES* and *COL1A1* are markers of the late stage of odontoblastic differentiation, regulated through the Wnt/β-catenin and BMP2/JNK pathways, while *ALPL* is a marker of the early stage of odontoblastic differentiation (*34*). *OMD*, a heterologous protein of Osteoadherin, belongs to the family of small leucine-rich proteoglycans and promotes the odontogenic differentiation (*35*). In contrast to previous studies indicating a predominance of unmyelinated nerve fibers over myelinated ones in the dental pulp (*36*), our findings suggest the opposite. In a healthy pulp, myelinated and unmyelinated cells account for 46.3% and 20.4% of the glial cell population, respectively. Additionally, odontoblast cells constitute 33.3% of the population, highlighting their significant role in dental pulp metabolism (Figure 5H).

### Progressive vascular responses to decay: Blood vessels early outward remodeling and arterialization followed by pathological angiogenesis and regression of the vascular network at the dentin-pulp interface

Dental decay progresses slowly, with early stages (Stages 1 and 2; SiSta classification) characterized by enamel demineralization and involvement of the outer and middle third of the dentin, while advanced stages (Stages 3 and 4; SiSta) progress to the inner third of the dentin and pulp tissue (Figure 3A). Despite early decay being physically distant from the pulp, we observed that pulp cells sense aggression already at Stage 1, triggering early inflammation and vascular remodeling. At Stage 1, arterialization is observed with the recruitment of α-SMA^+^ cells around capillaries in the capillary plexus (CP) on the decay-facing side of the pulp, contrasting with the absence of capillary arterialization on the healthy side (Figures 3B–3D). Early vascular responses include significant outward remodeling of capillaries already at stage 1, followed by venules at stage 2 and arterioles at stage 3 as the disease advances (Figure 3E). Increased blood flow, necessary to supply nutrients to activated odontoblasts and immune cells acting as first responders, leads to arterial-like pulsatile and high-pressure flow. This induces blood vessel differentiation such as capillary arterialization (*37, 38*). The increased relative expression of markers such as SELE, CXCL2, ICAM1, NFKBIA, CCL2, and VCAM marks endothelial cell (EC) activation (*39*). Arterialization is characterized by upregulating gene markers in mural cells, including *NOTCH3, MYL9, ACTA2, PDGFRB, MFGE8, RGS5, TAGLN, and BGN* (*40–42*). Several inflammatory mediators and neuropeptides, such as *NOS3, BDKRB2, HRH1, CXCL8, IL6R, IL1B, IL4R, IL7,* and *TACR1*, have their expression increased (Figures 4C–4E). NOS3, a marker of vasodilation (*43, 44*), is upregulated early and continues to increase progressively in venules and arterioles as the disease advances (Figure 4B). The bradykinin receptor B2, encoded by the *BDKRB2* gene, is a G-protein-coupled receptor for bradykinin, mediating biological effects such as inflammation and vasodilation (*45*). The histamine H1 receptor (*HRH1*) is a well-known inflammatory mediator, while interleukin-8 (*CXCL8*) is a potent inflammatory chemokine that promotes angiogenesis by activating the VEGF signaling pathway (*46, 47*). Additionally, tachykinin receptor 1 (*TACR1*), also known as the substance P receptor (*SPR*) or neurokinin 1 receptor (*NK1R*), is a G-protein-coupled receptor. It is found in both the central and peripheral nervous systems, functioning as a potent vasodilator (*48*). Our data show that in the early stages of the disease, the initial inflammatory response is associated with an immediate vascular response with capillary phenotypic variations.

**Figure 3.**
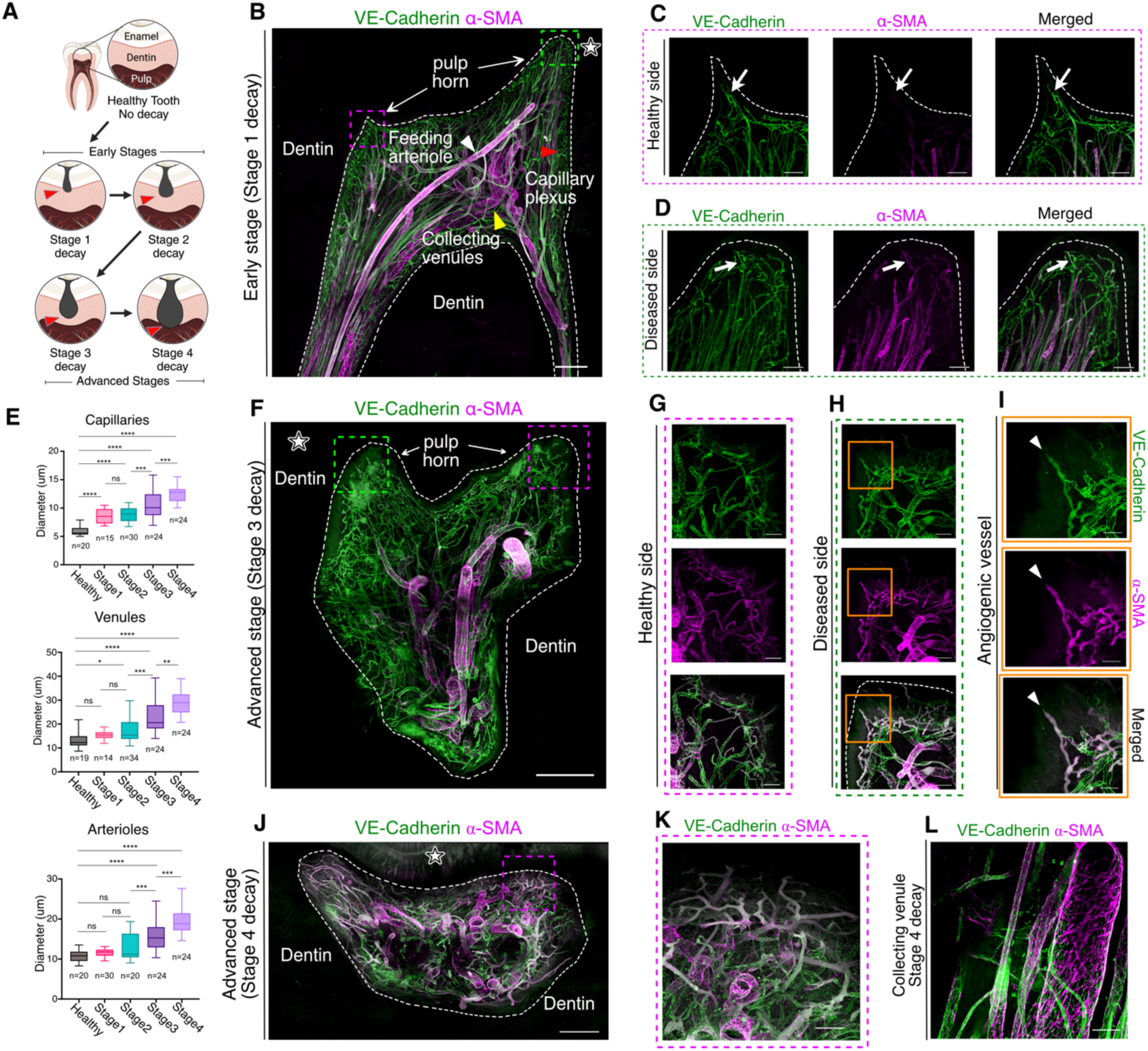
**Progressive vascular responses to decay: Blood vessels early outward remodeling and arterialization followed by pathological angiogenesis and regression of the vascular network at the dentin-pulp interface** (A) Schematic showing the different layers composing the tooth and the progression of the dental decay, from Healthy through Early to Advanced stages. (B) Maximum intensity projection (MIP) of a 486μm z-stack showing the vasculature of a mature dental pulp at an early stage (stage 1) of the disease. (C) MIP of 317μm z-stack of the pulp horn on the healthy side showing the capillary plexus. (white arrow) capillary without expression of α-SMA. (D) MIP of 308μm z-stack of the pulp horn on the diseased side showing the capillary plexus. (white arrow) capillary with expression of α-SMA.(E) Quantification of different vessel’s diameter increase (outward remodeling) on healthy dental pulps (n=6) and at various stages of their disease (n=21). (F) MIP of a 330μm z-stack showing the vasculature of a mature dental pulp at an advanced stage (stage 3) of the disease. (G) MIP of a 217μm z-stack showing the pulp horn on the healthy side showing non-perfused vessels. (H) MIP of a 338μm z-stack showing the pulp horn on the diseased side showing non-perfused vessel sprout and potential pathological angiogenesis.(I) Zoom on the vessel sprout (white arrowhead). (J) MIP of a 150μm z-stack showing the vasculature of a mature dental pulp at an advanced stage (stage 4) of the disease. (K) MIP of a 335μm z-stack showing non-functional and regressed capillary plexus. (L) MIP of a 125μm z-stack showing a heavily dilated collecting venule at stage 4 of the disease. Tissue clearing using a modified iDISCO protocol and staining for VE-Cadherin (green) and α-SMA (purple). Point scanning resonant confocal microscope with variable spectrum detector (Nikon AX R). Large images for (B, F, J). Plan Apo 10x λS OFN25 DIC N1 optic for (B, D, F). Plan apo Lambda 25XC Sil. optic for (C, E, G, H, I, J, K, L). Scale bars: 500µm for (B, F, J); 100µm for (C, D, G, H, K, L); 50µm for (I).

**Figure 4.**
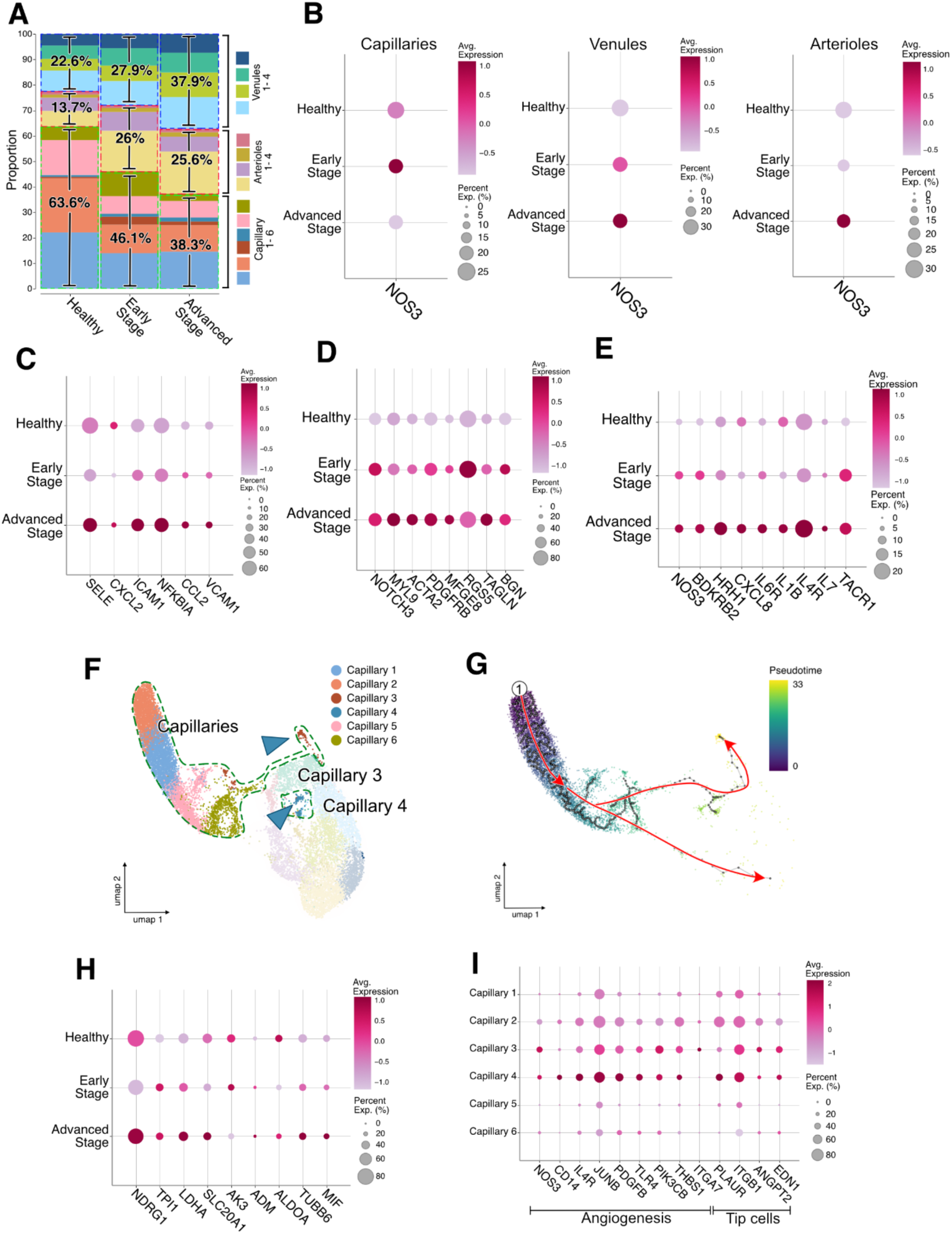
**Single-cell RNA Seq analysis of the endothelial sub-clusters of human dental pulps at various stage of their disease.** (A) Proportions of different endothelial sub-clusters according to their health states (healthy, early and advanced stage disease). (B) Dot plots showing a well-established marker (NOS3) associated with blood vessels vasodilation. (C-E) Dot plot showing 6-8 well-established markers associated with (C) ECs activation, (D) ECs arterialization and (E) early inflammation. (F) UMAP representation of 15,435 filtered and batch corrected healthy and diseased dental pulp ECs across 4 healthy pulps, 4 early stage and 4 advanced stage diseased pulps. Capillary sub-clusters 3 and 4 are of special interest (blue arrow heads). (G) UMAP plot of human ECs isolated from human dental pulps. The red lines, which were drawn manually, indicate the major trajectory flows. (H) Dot plot showing 9 well-established markers associated with tissue hypoxia at different states of the disease. (I) Dot plot showing 13 well-established markers associated with Angiogenesis and with tip cells.

**Figure 5.**
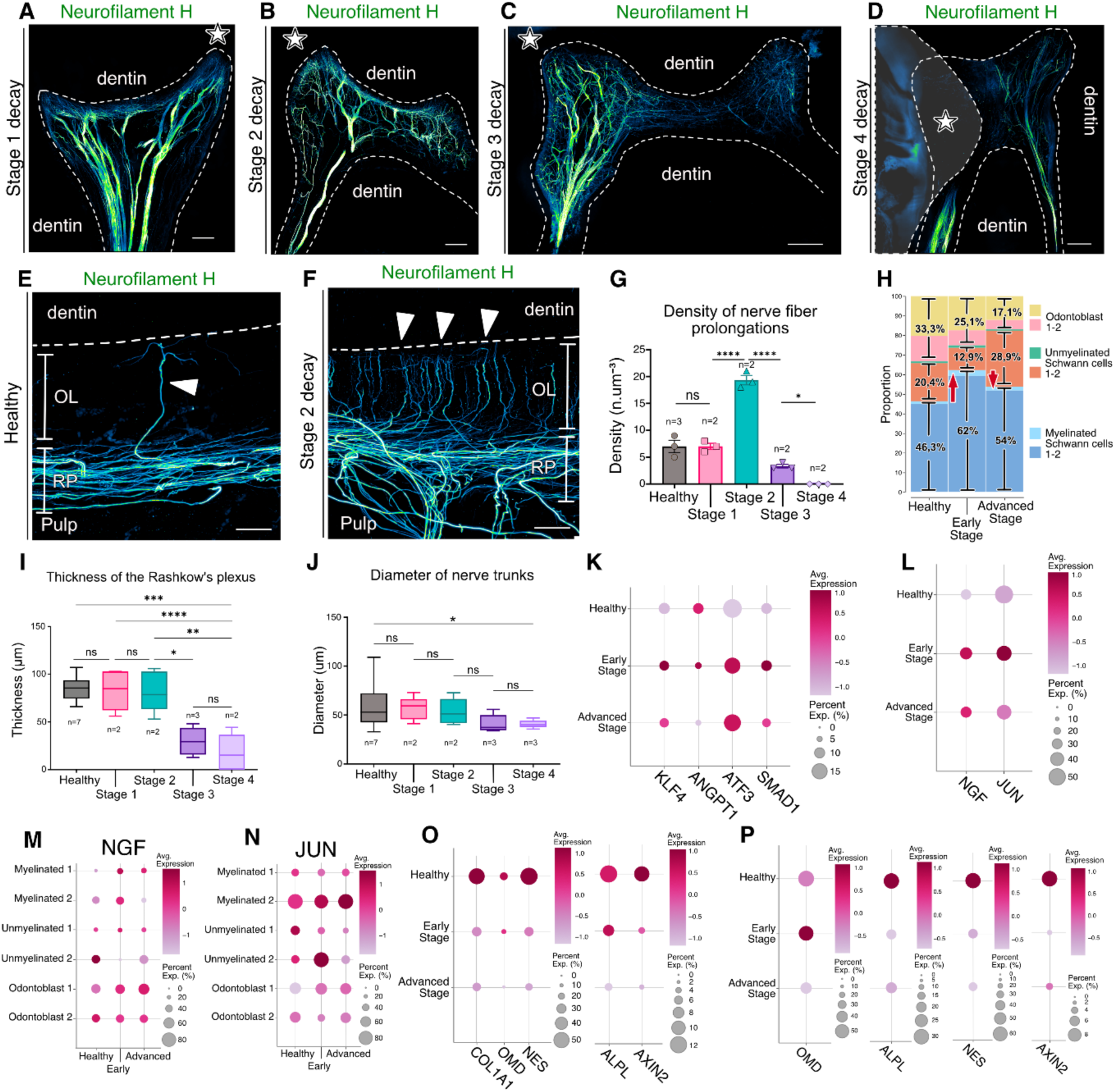
**Neurogenesis develops at the dentin-pulp interface in the early stages of the disease followed by Neural Regression at later stages** (A-D) Maximum intensity projection (MIP) of z-stack showing the neural network of a mature dental pulp at various stage of the disease. (white star) side where lays the decay. (A) stage 1 (233μm z-stack), (B) stage 2 (284μm z-stack), (C) stage 3 (293μm z-stack) and (D) stage 4 (291μm z-stack) of the disease. (E) MIP of a 140μm z-stacks in a healthy dental pulp. (F) MIP of a 205μm z-stacks in a diseased dental pulp at stage 2. (RP) Rashkow’s plexus; (OL) the odontoblastic layer. (G) Graph showing the density of nerve fibers prolongations (in n.μm-3) et different health states. (H) proportions of different sub-clusters at different health states. (I) Graph showing the thickness of the RP at different health states. (J) Graph showing the diameters of nerve trunks at different health states. (K-L) Dot plot showing 2-4 well-established markers associated with (K) nerve damage and (L) axon growth. (M) Dot plot showing the relative expression of NGF at different health states. (N) Dot plot showing the relative expression of JUN at different health states. (O) Dot plot showing 5 well-established markers for odontoblast cells at different health states. (A-F) Tissue clearing using a modified iDISCO protocol and staining with Neurofilament F (green) for the neural network. Point scanning resonant confocal microscope. Large images for (A-D). Plan Apo 10x λS OFN25 DIC N1 optic for (A-D). Plan apo Lambda 25XC Sil. optic for (E, F). Scale bars: 500µm for (A-D), 50µm for (E, F).

As the disease progresses, inflammation spreads within the pulp, leading to capillary regression and tissue hypoxia. Hypoxia is marked by increased expression of genes such as *NDRG1, TPI1, LDHA, SLC20A1, AK3, ADM, ALDOA, TUBB6*, and *MIF* (Figure 4H) (*49–51*). This correlated with the presence of vessel sprouting and angiogenesis, observed at Stage 3, where new blood vessels form to address oxygen deficiency. However, these vessels lack endothelial junctions and lumens, indicating a non-functional, pathological angiogenesis state which cannot resolve local hypoxia (Figures 3F–3I). Specific subclusters of capillaries, such as Capillary 3 and 4, expand during early and advanced stages of decay (Figure 4A). Trajectory analysis shows differentiation toward these angiogenic subclusters (Figures 4F–4G). Angiogenesis markers (*NOS3, CD14, IL4R, JUNB, PDGFB, TLR4, PIK3CB, THBS1, ITGA7*) (*52, 53*) and tip cells markers (*PLAUR, ITGB1, ANGPT2, EDN1*) (*54*) are significantly upregulated in late stages (Figure 4I). Pathological angiogenesis, vessel regression, tissue degradation, and outward remodeling extend throughout the pulp as the disease advances (Figures 3J–3L). These progressive vascular responses illustrate the transition from reparative (early stages) to pathological processes (late stages) in the dental pulp in response to tooth decay. Of importance, although we observe an extensive disorganization of the vascular network at later stages of the disease, we also observe remaining functional blood vessels under the fibrotic lesions and in the roots.

### Neurogenesis develops at the dentin-pulp interface in the early stages of the disease followed by Neural Regression at later stages

Progression of dental decay reveals increasing spatial disorganization of nerve fibers, alongside a reduction in nerve trunk diameters (Figures 5A–5D, 5J). In healthy pulps, the Raschkow’s Plexus (RP) resides beneath the odontoblastic layer (OL), with nerve fibers running parallel to the inner dentin surface. A limited number of fibers extend from the RP, cross the OL, and penetrate the dentin through dentinal tubules alongside odontoblastic prolongations. In Stage 2 decay, there is a notable increase in nerve fiber density across the OL (Figures 5E–5G) (*55*), potentially explaining heightened tooth sensitivity at this stage (*56, 57*). Myelinated nerve fibers, mainly present in the RP (*58*), transmit sensory signals. Single-cell analysis reveals an increase in the proportion of cells within Myelinated Schwann Cell Subcluster 1 at early stages (Figure 5H). Additionally, elevated expression of axon growth-associated gene markers, such as *NGF* (*59*) and *JUN* (*60*), is observed, particularly in Subcluster Myelinated Schwann Cells 1 (Figures 5L, 5M). Transitioning from Stage 2 to Stage 3, there is a significant thinning of the RP (Figure 5I). Early pulp inflammation triggers nerve inflammation, marked by increased expression of genes such as *KLF4* (*61*)*, ANGPT1* (*62*)*, ATF3* (*63*)*, and SMAD1* (*64*).

Odontoblastic cells, critical for tooth defense, are non-mitotic and naturally decline with age. Under aggression, they activate but are susceptible to death if the damage is severe. As decay progresses, a marked reduction in the relative expression of odontoblast-associated genes (*COL1A1, OMD, NES, ALPL, AXIN2*) is detected (Figure 5O) (*31–33, 65*), reflecting the detrimental impact of disease on the odontoblastic layer, necessary for dentin homeostasis (*66*).

### Early immune response followed by macrophage recruitment and activation as the diseases progresses

Single-cell analysis identified diverse clusters of blood and immune cells labeled with gene markers: *EBF1* for B-cells, *HLA*-*DRA* for dendritic cells (*67*), *CD14* for monocytes (*68*), *CD68* for macrophages (*69*), *PARP8* for NK cells (*70*), and *CD3E* for T-cells (*71*) (Figures 6A and 6D). The major immune cell populations are macrophages/monocytes/dendritic cells (37.4%), followed by B-cells (37%) and NK-cells & T-cells (25.6%) (Figure 6B). Macrophages were further categorized into subpopulations based on marker genes: M2 macrophages (*CD163, MRC1, CSF1R*) and M1 macrophages (*CD86, FCGR3A, RCGR2A, ITGAM*) (Figures 6C and 6E) (*72*). Inflammation, a complex response to harmful stimuli like bacteria and toxins, activates immune system components. Consistent with this, macrophage proportions and their relative gene expression levels increase as the disease progresses (Figure 6F). Activated macrophages, particularly the M2 subtype, play a pivotal role in crosstalk with fibroblasts. M2 macrophages secrete IL-10 and TGF-β, inducing fibroblast differentiation into myofibroblasts through macrophage-to-myofibroblast transition (MMT) (*73*). This process contributes to reactive tissue fibrosis (*73, 74*). Conversely, M1 macrophages are pro-inflammatory, accumulating around vascular injuries, and promoting endothelial cells (ECs) to upregulate vessel sprouting-associated genes, such as *ADAMTS-9*, *CTSS*, *CXCR4*, *EDNRB*, *KDR*, *NID1*, *NRCAM*, *PFKFB3*, *PLAUR*, and *UNC5B* (*75*).

**Figure 6.**
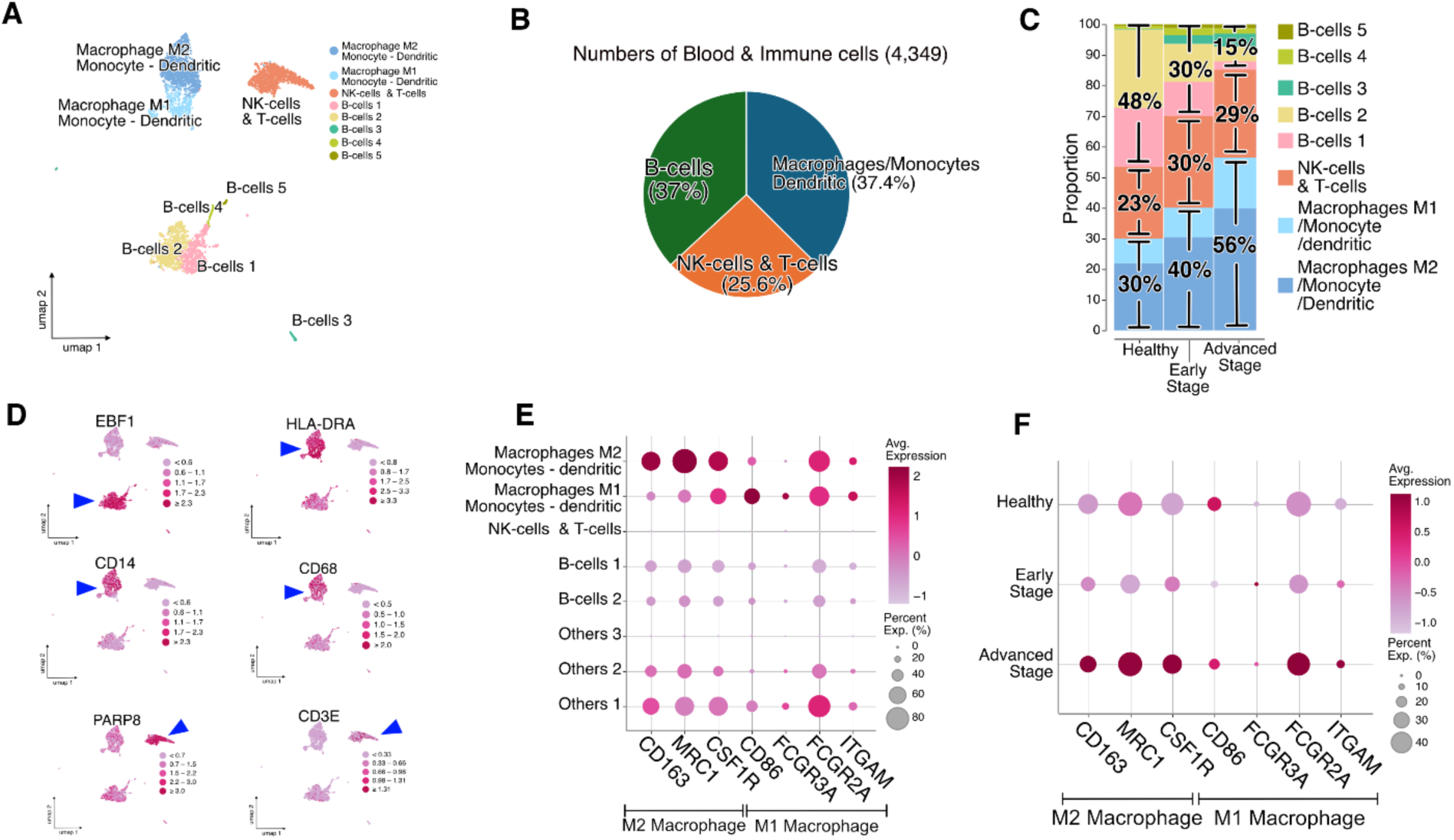
**Early immune response followed by macrophage recruitment and activation as the diseases progresses** (A) Uniform manifold approximation and projection (UMAP) representation of 4.349 filtered and batch corrected healthy and diseased dental pulp cells across 4 healthy pulps, 4 early stage and 4 advanced stage diseased pulps. (B) Proportions of each cell population. (C) Proportions of different immune cells sub-clusters according to their health states (healthy, early and advanced stage disease). (D) Relative expression of a well-established cell type marker for each cell population projected on UMAP plots. EBF1 for B-cells; HLA-DRA for dendritic cells; CD14 for Monocytes; CD68 for Macrophages; PARP8 for NK cells; CD3E for T-cells. (E) Dot plot showing 7 well-established markers associated with M1 and M2 Macrophages. (F) Dot plot showing 7 well-established markers associated with macrophages at different stages of the disease.

### Perivascular Progenitor Cell Differentiation into fibroblasts at the initiation of the disease is followed by myofibroblasts activation and reactive fibrosis in Advanced Disease Stages

In advanced stages of tooth decay, extensive tissue degradation is observed, with a necrotic zone directly beneath the decay site. Adjacent to this necrotic zone, we observe a fibrotic zone without any vascular and nervous structures but abundant α-SMA-expressing myofibroblasts (Figures 7A-C). Myofibroblast subclusters were identified using gene markers such as *ACTA2, TAGLN, AOC3, and MYH11* (Figures 7D-F) (*76, 77*), with their relative expression levels significantly increasing as the disease progresses. Additional subclusters were identified: fibroblasts (*VIM, CXCL14, LUM, C1S, DCN, FBLN2, POSTN, COL1A2, and CDH11*) (*78*), perivascular progenitor cells (*GLI1, CSPG4*) (*79, 80*) (Figures 7D and I), and mesenchymal progenitor cells (*MYC, ENG, NT5E, KLF4, and THY1*) (*81*) (Figures 7D and H). In healthy tissue, perivascular progenitor cells are prevalent; however, this cellular population disappears at the onset of tooth decay (Figures 7E and J). This indicates the early differentiation of Gli1^+^ perivascular progenitor cells into fibroblasts (subclusters 2, 3, and 4), whose numbers increase significantly during disease progression. This differentiation could be triggered by the early activation of the capillary network as part of the reparative response. Subsequently, these fibroblasts further differentiate into myofibroblasts under activation by M2 macrophage as the disease progresses (*73*). This differentiation pathway was confirmed by a trajectory analysis (Figure 7K).

**Figure 7.**
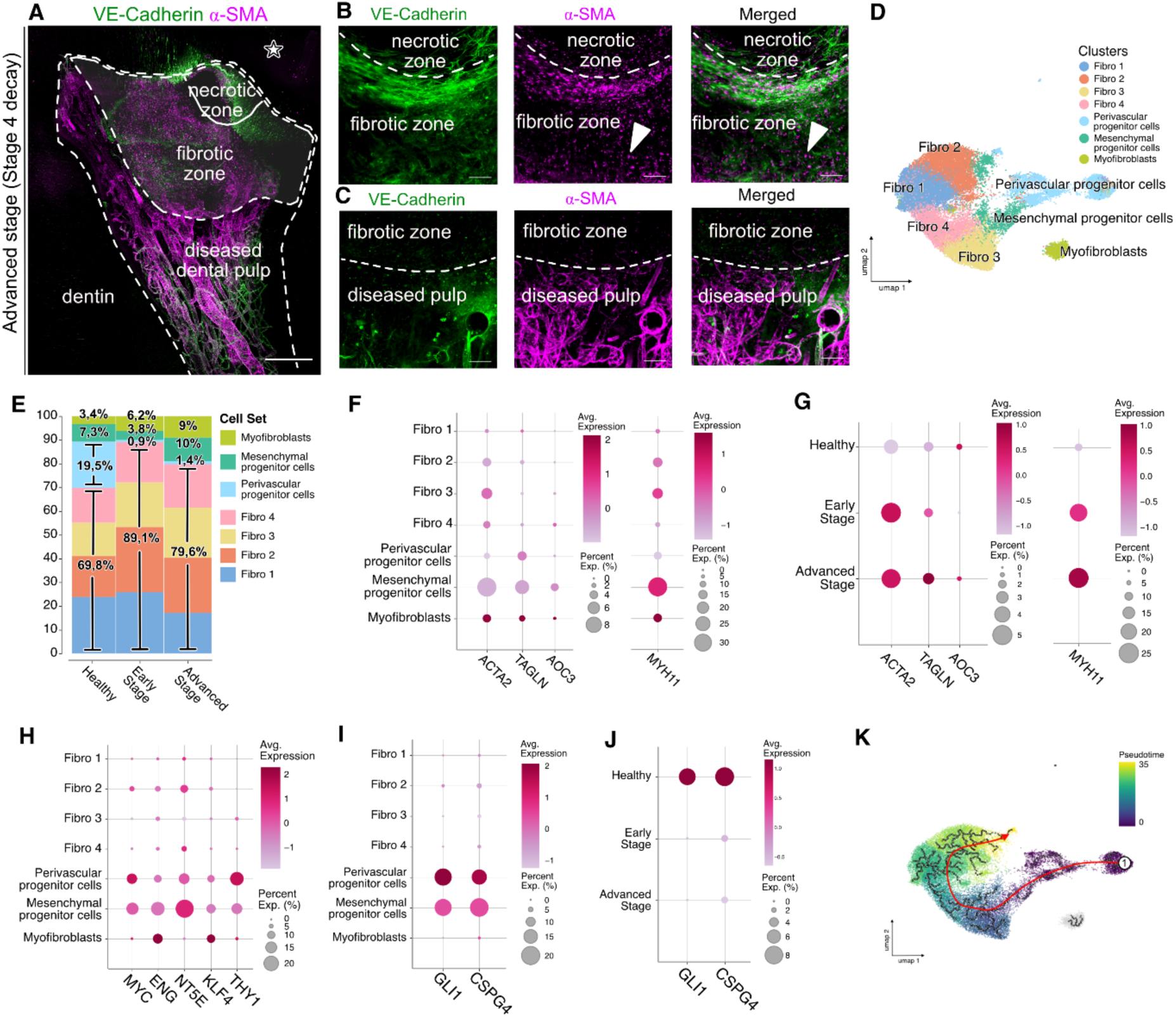
**Perivascular Progenitor Cell Differentiation into fibroblasts at the initiation of the disease is followed by myofibroblasts activation and reactive fibrosis in Advanced Disease Stages** (A) Maximum intensity projection (MIP) of a 152μm z-stack showing the presence of necrotic and fibrotic zone in a mature dental pulp at an advanced stage (stage 4) of the disease. (B) MIP of 181μm z-stack showing the transition between the necrotic and fibrotic zones. (C) MIP of 122μm z-stack showing the transition between the fibrotic zone and the adjacent diseased dental pulp. (D) Uniform manifold approximation and projection (UMAP) representation of 28.419 filtered and batch corrected healthy and diseased dental pulp fibroblast and stromal cells across 4 healthy pulps, 4 early stage and 4 advanced stage diseased pulps. (E) Proportions of different fibroblasts and stromal cells sub-clusters according to their health states (healthy, early and advanced stage disease). (F-G) Dot plot showing 4 well-established markers associated (F) with myofibroblasts. (G) with myofibroblasts at various health states. (H) Dot plot showing 5 well-established markers associated with progenitor stem cells. (I-J) Dot plot showing 2 well-established markers associated (I) with perivascular progenitor cells and (J) with perivascular progenitor cells at various health states. (K) UMAP plot of human fibroblast and stromal cells isolated from human dental pulps. The red lines, which were drawn manually, indicate the major trajectory flows. Tissue clearing using a modified iDISCO protocol and staining for VE-Cadherin (green) and α-SMA (purple). Point scanning resonant confocal microscope. Large images for (A). Plan apo Lambda 25XC Sil. optic for (A-C). Scale bars: 500µm for (A); 100µm for (B, C).

## Discussion

The tooth is a unique ectodermal organ composed of interconnected hard and soft tissues that interact dynamically at the dentin-pulp interface, forming the dentin-pulp complex (*82*). Our study aimed to elucidate the underappreciated impact of hard tissue (dentin) damage on distant soft tissue (dental pulp) during the natural course of the disease. By integrating high-resolution volumetric confocal imaging with in-depth single-cell analysis, we created the most comprehensive imaging and cellular atlas of healthy and diseased human teeth to date. Furthermore, our findings have significant global implications, as they reveal that dental pulp responses mirror those observed in other tissues under pathological conditions.

Our panoramic approach highlights the intricate interplay between various systems within the dental pulp throughout disease progression, with the vasculature playing a pivotal role since most components involved in the inflammatory response transit through it (*83*). Early in the disease, bacterial invasion of the outer dentin layer is detected by specialized defense cells within the pulp, called odontoblasts, specifically through their odontoblastic processes. An arterialization of capillaries occurs at the localized inflamed site, recruiting alpha-SMA^+^ cells driven by chemoattractant-VEGF produced by stress-activated endothelial cells (*84*). Capillary lumen enlargement (outward remodeling) occurs as a mechanical response to the increased blood flow at the inflamed site. Similarly, in the early stages of atherosclerosis, a radial expansion of vessels help preserve lumen size (*85, 86*). Appropriate inflammation acts as a self-protective mechanism, promoting nerve and axon regeneration and myelin formation (*87*). Our data demonstrates that neurogenesis, through nerve-endings sprouting, increases inside the odontoblastic layer at an early stage, explaining the increased tooth sensitivity the patient feels as peripheral inflammatory mediators induce peripheral nerve sensitization (*88*). The myelinated Schwann cells exhibit increased synthesis of Nerve Growth Factor and pose as the primary mediators of this early neurogenesis. Additionally, Gli1^+^ perivascular progenitor cells rapidly differentiate into fibroblasts, contributing to the pool of activated fibroblasts essential for tissue repair (*89*). At these early stages of the disease (stages 1&2), the various remodeling processes contribute to tissue repair and aiming for a return to homeostasis. Our data have an important clinical implication. At this stage, early lesions can be stabilized through remineralization if appropriate measures are taken, such as implementing a dental hygiene program, modifying the diet, and ensuring proper fluoride intake. If conservative treatment, such as placing a dental filling, is required, it will remove the demineralized tissue along with the microorganisms responsible for pulp inflammation. Our findings suggest that an early intervention on the affected mineralized tissues is essential to allow a natural repair of the underlying dental pulp. Further research is needed to investigate the pulp’s return to homeostasis.

Our data highlights the importance of containing the inflammatory process as excessive and sustained inflammation elicits irreversible remodeling. As decay advances toward the dental pulp and the inflammation becomes chronic, an increasing number of mononuclear leukocytes (monocytes) are recruited to the inflamed and hypoxic tissue. These cells differentiate into macrophages, promoting the formation of new blood vessels, called angiogenesis. As the formation of these de novo vessels originate from preexisting capillary bed, this phenomenon refers to “sprouting angiogenesis”. Tissue hypoxia, as seen in various diseases such as cancer or atherosclerosis, plays the role of angiogenic switch (*90*). Our study shows a regression of vessels in the capillary plexus with the progression of the disease, leading to a lack of oxygen necessary for the odontoblast’s activity. However, our data also shows that newly formed vessels are often non-perfused and non-functional due to the absence of endothelial junctions and lumens, constituting pathological angiogenesis. In other part of the body, the latter is also observed and can play a supportive role for pathological processes such as tumor angiogenesis, atherosclerosis neovascularization or age-related macular degeneration. It can be present in systemic diseases such as coronary artery disease or peripheral arterial disease (PAD), when the angiogenic response is insufficient to restore an adequate and functional vasculature (*91*). Persistent angiogenesis exacerbates macrophage infiltration and precipitates tissue damage (*92*). Chronic inflammation also increases peripheral nerve damage, releasing additional inflammatory mediators and perpetuating the inflammatory cycle (*88*). Simultaneously, activated fibroblasts increasingly differentiate into alpha-smooth muscle actin^+^ myofibroblasts, producing collagen and other ECM proteins, resulting in tissue fibrosis and degradation, as observed in many other tissues such as the heart or the lungs. At these advanced stages, merely removing the affected hard tissues, as is commonly done in clinical practice today, is insufficient to stop the progression of tissue remodeling and degradation. This ultimately leads to the need for complete dental pulp removal from the coronal and the root portions, replacing it with synthetic materials. This procedure, known as pulpectomy or root canal therapy, is commonly used to manage mature teeth with carious pulp involvement (*93*). In contrast, a pulpotomy involves only the partial removal of the dental pulp, limited to its coronal section, with the remaining tissue covered by a calcium silicate-based cement. It is the recommended procedure for primary or immature teeth to preserve pulp vitality until root formation is complete, as well as for mature teeth with irreversible pulpitis (*93*). The successful outcome of this treatment relies solely on clinical and subjective criteria, such as the absence of pain and the lack of radiographic signs of peri-apical lesions (*93*). In the event of a pulpectomy, a whole pulp regeneration is a promising future therapy, not yet achievable. Our findings indicate that interventions at advanced disease stages require a multimodal approach. Restoring the affected hard tissues with dental materials should be complemented by strategies aimed at arresting and reversing dental pulp damage. These may include novel treatments such as angiogenesis inhibitors, antifibrotic drugs, or innovative regenerative therapies. These novel treatments associated with procedures such as a complete pulpotomy, which is the removal of the dental pulp in the coronal portion until the entry of the roots, or a partial pulpotomy, which is a partial removal (2-3 mm) of the coronal pulp (*94*), could become a new standard of care. Indeed, root canal therapies totally destroying the pulp tissue seem unnecessary and prevent the possibility for future pulp regeneration therapies. Further research is needed to investigate ways to promote pulp regeneration from the remaining living tissue, unaffected by fibrosis.

Contrary to the current belief that odontoblast cells are activated in response to aggression and that mesenchymal cells differentiate into neo-odontoblasts (odontoblast-like cells) to produce tertiary protective dentin (*34, 95*), our data do not indicate the activation of most genes associated with odontoblastic differentiation (*ALP, NES, AXIN2*). These findings challenge and redefine the role of odontoblasts in the dental pulp defense mechanism.

Untreated advanced dental decay is often linked to intensified and exacerbated pain. However, this clinical observation contrasts with our data, which show an increase in nerve fiber density (neurogenesis) at early stages, followed by a gradual decline in later stages. These findings suggest the need to rethink current concepts of dental pain (*95*). Pain in the early stages is likely associated with nerve endings in Raschkow’s plexus, while pain in later stages may be linked to the autonomic nervous system and its connections to blood vessels.

Unlike developing and immature dental pulp, which contains stem cells known as human dental pulp stem cells (hDPSCs), healthy mature dental pulp contains Gli+ perivascular progenitor cells and mesenchymal progenitor cells. Our findings reveal a drastic decline in Gli+ perivascular progenitor cells as soon as the disease begins, emphasizing the critical role of the vasculature in the immediate tissue response to inflammation. Additionally, it has been previously suggested that Gli1^+^ cells proliferate and differentiate into myofibroblast following injury to organs such as the heart, liver, lung or kidney (*79*). Our study revealed that disease progression to advanced stages leads to the establishment of a fibrotic condition, similar to degenerative mechanisms present in other human organs.

In conclusion, our findings emphasize that dental decay is a slow-progressing disease characterized by a continuum of transitional states within the pulp’s cellular environment. These states involve multiple system interplays and a delicate balance between reversible reparative processes (e.g., outward remodeling and arterialization) and irreversible pathological mechanisms (e.g., pathological angiogenesis and fibrosis). The implications extend beyond dental sciences. In addition to establishing new standards of care for the dental profession, understanding these processes in the tooth will provide valuable insights into systemic conditions and offer a platform for testing innovative therapeutics.

## Limitations of the study

While this study provides for the first time a combined imaging and cellular insights into the biological response of the dental pulp to decay on untreated teeth, a limitation should be acknowledged. The healthy and diseased samples analyzed in this research were not always obtained from the same patient, which introduces variability due to individual genetic and physiological differences. An uncompleted tissue dissociation can also result in variation in cell proportion in the scRNA seq analysis.

## Supplemental information

Raw (FASTQ files) and processed data from the single-cell experiment have been deposed at ArrayExpress (functional genomic data collection) and are publicly available as of the date of publication (ArrayExpress accession E-MTAB-14896).

## Acknowledgments

The authors thank Prof. Laurence Evrard, Mrs. Fairouz Ben Abdeloualed and all the clinical contributors from the Dentistry and Stomatology department at the Erasme University Hospital (HUB -Erasme site, Brussels, Belgium) for providing the human tissue samples. The authors also thank Mrs. Emilie Calonne from the laboratory of cancer epigenetic (Prof. François Fuks, ULB, Brussels, Belgium) for her technical support. They also thank Prof. Cédric Blanpain (Laboratory of Stem Cells and Cancer, ULB, Brussels, Belgium) for his critical insights. This project was financially supported by funding from the Foundation ULB and a Carte Blanche funding from ULB.

## Author contributions

H.H. and N.B. conceptualized and designed the project; H.H., C.G., V.V., E.G., S.S. and N.B. performed data analysis and interpretation of the data; H.H., S.K., A.V., I.T., O.K., H.B., D.S. performed the experiments; V.V. and E.G. provided the resources for the study; N.B. acquired fundings; H.H. and N.B. wrote the manuscript with all authors contributing to provide feedback, and gave their final approval for the work.

## Methods

### Key resources table

| REAGENT or RESOURCE | SOURCE | IDENTIFIER |
| --- | --- | --- |
| **Antibodies** |  |  |
| Polyclonal Goat IgG Human VE-Cadherin Antibody | R&D Systems, Inc. | AF938, RRID: AB_355726 |
| Mouse monoclonal Anti-α Smooth Muscle Actin | Sigma-Aldrich | Cat# A2547, RRID: AB_476701, Cat# A5228, RRID: AB_262054 |
| Neurofilament-H Rabbit monoclonal Ab | Cell signaling | Cat #30564 |
| Nestin Monoclonal Antibody | Thermo Fisher Scientific | Cat# 14-9843-80, RRID: AB_1548838 |
| Nestin Antibody | Cell signaling | Cat#10959 |
| UCHL1/PGP9.5 antibody | Proteintech | Cat# 14730-1-AP, RRID: AB_2210497 |
| Podoplanin antibody | Proteintech | Cat# 67432-1-Ig RRID: AB_2882670 |
| Human LYVE-1 Antibody | R&D Systems, Inc. | AF2089, RRID: AB_355144 |
| PROX1 Polyclonal antibody | Proteintech | Cat# 11067-2-AP, RRID: AB_2268804 |
| Donkey anti-Goat IgG (H+L) Highly Cross-Adsorbed Secondary Antibody, Alexa Fluor™ Plus 555 | Thermo Fisher Scientific | Cat# A32816, RRID: AB_2762839 |
| Donkey Anti-Mouse IgG H&L (Alexa Fluor® 647) | Abcam | Cat# ab150107, RRID: AB_2890037 |
| Donkey anti-Rabbit IgG (H+L) Highly Cross-Adsorbed Secondary Antibody, Alexa Fluor™ Plus 555 | Thermo Fisher Scientific | Cat# A32794, RRID: AB_2762834 |
| Goat anti-Mouse IgG (H+L) Highly Cross-Adsorbed Secondary Antibody, Alexa Fluor™ Plus 555 | Thermo Fisher Scientific | Cat# A32727, RRID: AB_2633276 |
| Donkey anti-Rabbit IgG (H+L) Highly Cross-Adsorbed Secondary Antibody, Alexa Fluor™ Plus 647 | Thermo Fisher Scientific | Cat# A32795, RRID: AB_2762835 |
| Donkey anti-Mouse IgG (H+L) Highly Cross-Adsorbed Secondary Antibody, Alexa Fluor™ Plus 647 | Thermo Fisher Scientific | Cat# A32787, RRID: AB_2762830 |
| **Biological samples** |  |  |
| Healthy and diseased adult human teeth | The Dentistry and Stomatology biobank at the Erasme University Hospital (HUB - Erasme site) (AFMPS number: BB190032) | https://www.erasme.be/fr/enseignement-recherche/comite-d-ethique/biobanques-cbeu/les-biobanques-sur-le-campus-erasme |
| **Chemicals, peptides, and recombinant proteins** |  |  |
| Ethylenediaminetetraacetic acid disodium salt dihydrate | Sigma-Aldrich | ED2SS-1KG |
| Methanol | Sigma-Aldrich | 179957-1L |
| Hydrogen peroxide solution 30% | Sigma-Aldrich | 216763-100ML |
| DMSO | Sigma-Aldrich | D128-4 |
| Triton X-100 | Sigma-Aldrich | X100-500ML |
| Tetrahydrofuran THF | Sigma-Aldrich | 186562 |
| Dichloromethane (DCM) | Sigma-Aldrich | 270997 |
| Dibenzylether (DBE) | Sigma-Aldrich | 108014-1KG |
| Nunc Glass base dish 27mm | ThermoFisher | 150682 |
| Intercept® (PBS) Blocking Buffer | LI-COR Biosciences - GmbH | 927-70001 |
| Collagenase Type 4 | Worthington Biochemical Corp. | LS004210 |
| Hank's Balanced Salt Solution | Sigma-Aldrich | H9394 |
| RPMI 1640 Medium, no glutamine, no phenol red 500 mL | ThermoFisher | 32404014 |
| TrypLE™ Express Enzyme (1X), phenol red | ThermoFisher | 12605010 |
| Cell strainer | VWR | 732-2758 |
| Antibiotic Antimycotic Solution (100×), Stabilized | Sigma-Aldrich | A5955-20ML |
| KAPA Pure Beads (5ml) | Roche | KK8000 (07983271001) |
| Critical commercial assays |  |  |
| Evercode™ Cell Fixation v2 | Parse biosciences | Ref# ECF2101 |
| Evercode™ WT v2 | Parse biosciences | Ref# ECW02130 |
| UDI Plate - WT | Parse biosciences | Ref# UDI1001 |
| Qubit™ dsDNA HS Assay Kit | ThermoFisher | Q32851 |
| Bioanalyzer High Sensitivity DNA Analysis | Aligent | 5067-4626; 5067-4627 |
| Software and algorithms |  |  |
| NIS-Elements | NIKON CORPORATION | https://www.microscope.healthcare.nikon.com/en_EU/products/software/nis-elements |
| GraphPad Prism 9 | GraphPad Software | https://www.graphpad.com/updates/prism-900-release-notes |
| Trailmaker™ | Parse biosciences | https://www.parsebiosciences.com/data-analysis/ |
| Other |  |  |
| Isomet Low Speed® | Buehler | https://www.buehler.com/products/sectioning/precision-cutters/isomet-low-speed-precision-cutter/ |
| Scanning point resonant confocal microscope with a variable spectrum detector (Nikon AX R) | Nikon | https://www.microscope.healthcare.nikon.com/en_EU/products/confocal-microscopes/ax |
| Confocal microscope with a spinning disk (X-Light V3, Crest Optics), | Nikon | https://www.microscope.healthcare.nikon.com/en_EU/products/confocal-microscopes/crest-x-light-series/crest-x-light-v3 |
| 10x immersion objective (N/A 0.5, WD 5.5 mm; Nikon Plan Apo) | Nikon | https://www.microscope.healthcare.nikon.com/en_EU/products/optics/selector |
| Plan Apo 10x λS OFN25 DIC N1 optics | Nikon | https://www.microscope.healthcare.nikon.com/en_EU/products/optics/selector |
| Plan Apo Lambda 25xC Sil | Nikon | https://www.microscope.healthcare.nikon.com/en_EU/products/optics/selector |
| Qubit™ Flex Fluorometer | ThermoFisher | https://www.thermofisher.com/be/en/home/industrial/spectroscopy-elemental-isotope-analysis/molecular-spectroscopy/fluorometers/qubit.html |
| 2100 Bioanalyzer | Agilent | https://www.agilent.com/en/product/automated-electrophoresis/bioanalyzer-systems/bioanalyzer-instrument |

### Experimental Model and Study Participant Details

### Human patient tissue samples

Healthy and diseased human teeth were obtained from the Dentistry and Stomatology biobank at the Erasme University Hospital (HUB - Erasme site) (AFMPS number: BB190032). The samples were classified as residual human body material and were collected from patients attending the Outpatient Dentistry Department and the One-Day Clinic at the Erasme University Hospital (HUB - Erasme site, Brussels, Belgium). This study received approval from the Erasme Hospital-Faculty Ethics Committee – ULB (Reference number P2021/146). All samples were anonymized to ensure patient identities were not accessible. The collection included healthy and diseased teeth from patients of both sexes, aged between 18 and 50 years, and representing various ethnicities (Table S1).

The SiSta (Site & Stage) classification was used to determine the degree of progression in diseased teeth. The teeth were categorized into five groups:

1. Stage 0: representing a healthy tooth or a tooth with decay strictly limited to the enamel layer.
2. Stage 1: representing a tooth with decay limited to the external third of the dentin.
3. Stage 2: representing a tooth with decay limited to the middle third of the dentin.
4. Stage 3: representing a tooth with decay extending to the internal third of the dentin; and
5. Stage 4: representing a tooth with decay extending to the internal third and involving the dental pulp.

### Imaging experimental protocol

#### Human patient tissue acquisition for imaging

Samples from healthy and diseased teeth were acquired from the Dentistry and Stomatology biobank (Erasme University Hospital, HUB - Erasme site, Brussels, Belgium). The samples were fixed in a 4% formaldehyde solution immediately after extraction and stored at 4°C for 7 to 14 days. The teeth were sectioned into 1 mm-thick slices in the mesiodistal direction for healthy teeth and along the direction of decay for diseased teeth, using a gravity-fed precision sectioning machine (Isomet Low Speed®, Buehler, Germany) equipped with a precision diamond blade (IsoMet® Diamond Wafering Blades, Buehler [Arbor Size: 0.5]). Sections containing both hard and soft tissues were preserved in Phosphate-Buffered Saline (PBS) solution at 4°C.

#### iDISCO clearing protocol

The original iDISCO protocol(*9*) was modified to suit our samples. Sectioned samples were first decalcified in ethylenediaminetetraacetic acid (EDTA) solution (0.5 M, pH 7.0), which was renewed daily for a total of 7 days and kept on a shaker at 37°C. They were washed in PBS solution for 1 day at room temperature. The samples were dehydrated at room temperature in methanol solutions diluted in PBS, as follows: 50% methanol for 1 hour, 80% methanol for 1 hour, and 100% methanol twice for 1 hour each.

The samples were bleached overnight at 4°C in a solution of 5% H_2_O_2_ in 20% DMSO/methanol, followed by washing steps at room temperature in 100% methanol twice for 1 hour and in 20% DMSO/methanol twice for 1 hour. The samples were rehydrated at room temperature as follows: 80% methanol for 1 hour, 50% methanol for 1 hour, and PBS twice for 1 hour each. This was followed by a blocking step at room temperature twice for 1 hour in PTx2 solution (PBS/0.2% Triton X-100).

The immunostaining step began with overnight permeabilization at 37°C in PTwH solution (1× PBS/0.2% Triton X-100/20% DMSO/0.3 M glycine). The samples were then blocked in Blocking Buffer solution (Intercept® Blocking Buffer, Li-Cor®) containing 0.2% Triton X-100 (Sigma-Aldrich) for 2 days and subsequently washed at room temperature twice for 1 hour in PBS.

The samples were then incubated in the blocking buffer solution containing 0.2% Triton X-100 and a primary antibody diluted at 1:1000 (1‰) on a shaker at 37°C for 3 days, followed by washing in PBS solution for 1 day at room temperature. From this stage onward, the samples were protected from light using aluminum foil. The samples were then incubated in the blocking buffer solution containing 0.2% Triton X-100 and a secondary antibody diluted at 1:1000 (1‰) on a shaker at 37°C for 3 days, followed by washing in PBS solution for 1 day at room temperature.

The samples were dehydrated overnight at room temperature in 50% v/v tetrahydrofuran/dH2O (THF) solution, then for 1 hour in 80% v/v THF/dH2O solution, and twice for 1 hour in 100% v/v THF solution. The samples were then incubated at room temperature in dichloromethane (DCM) until they sank to the bottom of the vial. Finally, the samples were cleared at room temperature in dibenzyl ether (DBE) until fully cleared and stored in the same solution.

All the steps were performed on a shaker.

### Microscopy and image analysis

Fluorescent images were acquired using two different microscopes. First, a scanning point resonant confocal microscope with a variable spectrum detector (Nikon AX R), equipped with various optics such as a Plan Apo 10x λS OFN25 DIC N1 and a Plan Apo Lambda 25xC Sil, was used to visualize the markings. Samples were mounted on Nunc™ Glass bottom dishes (Thermo Fisher Scientific, Massachusetts, USA) along with the storage solution. Second, a confocal microscope (Nikon) with a spinning disk (X-Light V3, Crest Optics), connected to a camera (Photometrics Prime BSI), and equipped with a 10x immersion objective (N/A 0.5, WD 5.5 mm; Nikon Plan Apo), was also used to visualize the markings. Samples were mounted on Lab Glass Petri Dishes along with the storage solution. Z-stacks of various steps and various depth (up to 600µm) were acquired and processed using NIS software. The images were subsequently processed into Maximum intensity projections (MIP) on which quantifications were performed.

### Statistical analysis

Statistical analysis was performed using the GraphPad Prism 9 software (GraphPad Software, Boston, USA). Data are expressed as mean ± SEM and distribution is displayed within the graphs. Various tests were used to calculate the P values and are indicated in the figure legends.

### Human patient tissue acquisition and tissues dissociation for single cell

Samples from healthy and diseased teeth were collected from patients attending the Outpatient Dentistry Department and the One-Day Clinic at the Erasme University Hospital (HUB - Erasme site, Brussels, Belgium). Immediately after extraction, the samples were placed in Hank’s Balanced Salt Solution (HBSS, Sigma-Aldrich, St.Louis, USA) containing 1% Penicillin/Streptomycin/Amphotericin B solution (A5955-100ML, Sigma-Aldrich, St.Louis, USA) and kept on ice. All samples were processed and dissociated within 60 minutes of extraction.

Fresh collagenase solution for tissue dissociation was prepared by resuspending 40 mg of Collagenase Type 4 (Worthington Biochemical, NJ, USA) and 4 mg of calcium chloride in 10 mL of prewarmed TrypLE™ Express Enzyme 1X phenol red (Thermo Fisher Scientific, Massachusetts, USA) and maintained at 37°C. Dissection forceps and scissors were disinfected using a VIRKON S solution (Germineo, Onet Le Chateau, FRANCE). Petri dishes were filled with sterile, cold RPMI solution containing 1% Penicillin/Streptomycin/Amphotericin B.

To process the tooth, it was first held using a tooth holder, and the periodontium was scraped off using a surgical blade. The surface of the tooth was carefully wiped with 70% ethanol. The tooth was then cracked using a hammer, ensuring minimal compression to avoid tissue damage. The cracked fragments were placed into a petri dish containing cold RPMI solution and kept on ice.

Under a stereomicroscope and using forceps, the fragments were carefully separated to expose the dental pulp. The pulp was then extracted from the pulp chamber, transferred to a fresh petri dish filled with cold RPMI/antibiotic-antimycotic solution, and kept on ice. The dental pulp tissues were finely chopped into small pieces using fine scissors and immediately resuspended in a sterile 15 mL Falcon tube containing 3 mL of prewarmed collagenase solution.

The tubes were incubated at 37°C for 50 minutes, with manual shaking performed every 10 minutes. After incubation, the tubes were placed on ice, and the tissue was manually triturated by pipetting up and down, first with a slightly cut P1000 tip and then with a normal P1000 tip, for 3 minutes until complete tissue disaggregation.

The cell suspension was filtered through a 40 µm cell strainer into a new 15 mL tube, rinsing the filter with 1 mL of RPMI solution. The tube was centrifuged at 500g for 10 minutes in a swinging bucket rotor. The supernatant was carefully decanted, and the pellet was resuspended in 375 μL of cold Cell Prefixation Buffer (Parse Fixation Kit v2, Parse Biosciences, Seattle, USA).

Next, 125 μL of cold Cell Fixation Solution (ECF2101 Evercode™ Cell Fixation v2, Parse Biosciences, Seattle, USA) was added to the suspension, mixed by pipetting three times with a P1000, and incubated on ice for 10 minutes. Then, 40 μL of cold Cell Permeabilization Solution (ECF2101 Evercode™ Cell Fixation v2, Parse

Biosciences, Seattle, USA) was added, mixed, and incubated on ice for 3 minutes. Finally, 500 μL of cold Cell Neutralization Buffer (ECF2101 Evercode™ Cell Fixation v2, Parse Biosciences, Seattle, USA) was added, and the tube was centrifuged at 500 × g for 10 minutes in a swinging bucket rotor.

The supernatant was carefully removed and discarded, and the pellet was resuspended in 75 μL of cold Cell Buffer (ECF2101 Evercode™ Cell Fixation v2, Parse Biosciences, Seattle, USA) and transferred to a 1.5 mL tube. A 10 μL aliquot of the cell suspension was taken for cell counting. This aliquot was mixed with 40 μL of 0.4% trypan blue, and the cells were counted using a Neubauer chamber.

To preserve the sample, 3.75 μL of DMSO (ECF2101 Evercode™ Cell Fixation v2, Parse Biosciences, Seattle, USA) was added, and the sample was stored in a Mr. Frosty freezing container at −80°C for up to 6 months, until further processing for the single-cell protocol.

### Single-cell Library Generation and Sequencing

Single-cell sublibraries were generated following the Evercode™ WT v2 user manual v2.3 (ECW02130 Evercode™ WT v2, Parse Biosciences, Seattle, USA). Samples were thawed, counted, and recorded into the Sample Loading Table provided. Based on the Sample Loading Table values, samples were diluted with Dilution Buffer and loaded into the Round 1 Plate. During this step, an in-situ reverse transcription reaction was performed, and well-specific barcodes were added. The cells were then pooled, centrifuged, and resuspended.

The pooled cells were mixed with the Ligation Master Mix and transferred to the Round 2 Plate, where an in-situ ligation reaction attached a second well-specific barcode to the 3′ end of the cDNA. After pooling and straining, Round 3 Ligation Enzyme was added to the sample, which was then loaded into the Round 3 Plate. During this step, a second in-situ ligation reaction introduced a third well-specific barcode, the Illumina TruSeq R2 sequence, and a biotin to the cDNA. The sample was subsequently pooled, strained, centrifuged, washed, and resuspended in Dilution Buffer.

The cells were counted and allocated into sublibraries following the guidelines in the Sublibrary Generation Table. These sublibraries were lysed and stored at −80°C until further processing. The barcoded cDNA was then captured on streptavidin-coated magnetic Binder Beads and washed to remove cellular debris. After an additional wash, a template switching reaction was performed on the captured cDNA, adding a 5′ adaptor. The captured cDNA was further washed and amplified using primers targeting the template switching (TS) sequence and the Illumina TruSeq R2 sequence.

The amplified cDNA was purified using a 0.8x SPRI bead cleanup. Its concentration and size distribution were assessed using fluorescent dyes (Qubit™ dsDNA HS Assay Kit) and capillary electrophoresis (High Sensitivity DNA Kit on the Agilent Bioanalyzer System). The barcoded and amplified cDNA was subsequently fragmented, end-repaired, and A-tailed in a single reaction. Fragmented and A-tailed DNA underwent size selection with sequential 0.6x and 0.8x SPRI bead cleanups. Adaptors containing the Illumina TruSeq R2 sequence were ligated to the 5′ end of the fragmented DNA, followed by another 0.8× SPRI bead cleanup.

Adaptor-ligated DNA was PCR-amplified using Illumina TruSeq R1 and R2 primers. This indexing PCR created sequencing libraries while adding i5/i7 unique dual indices (UDIs) as a fourth layer of cell barcoding. The sequencing libraries were size selected using a double-sided SPRI bead cleanup. Concentration and size distribution were measured again using fluorescent dyes (Qubit™ dsDNA HS Assay Kit) and capillary electrophoresis (High Sensitivity DNA Kit on the Agilent Bioanalyzer System). Finally, the sublibraries were stored at 4°C for up to 48 hours or at −20°C for up to 3 months. Sublibraries were submitted for sequencing with the NovaSeq X Plus Series (PE150, Novogene UK, Cambridge, UK).

### Data Processing and Analysis

Sequenced data were processed using Trailmaker™ (Parse Biosciences, Seattle, USA). FASTQ files were first analyzed through the Parse Biosciences pipeline to generate project-specific Insight modules for further analysis in Trailmaker™.

Data processing involved six key steps:

1. Cell Size Distribution Filter: Filtering cells based on size distribution.
2. Mitochondrial Content Filter: Filtering cells with high mitochondrial transcript content.
3. Number of Genes vs. Transcripts Filter: Filtering cells based on the number of detected genes and transcripts.
4. Doublet Filter: Removing doublets from the dataset.
5. Data Integration: Data were integrated using the Harmony method.
6. Configure Embedding: Nonlinear dimensionality reduction techniques, such as Uniform Manifold Approximation and Projection (UMAP), were applied to visualize clusters or groups of data points and their relative proximities.

Clustering was performed using the Louvain clustering method. The integrated data were LogNormalized before embedding.

---

**Figure S1.**
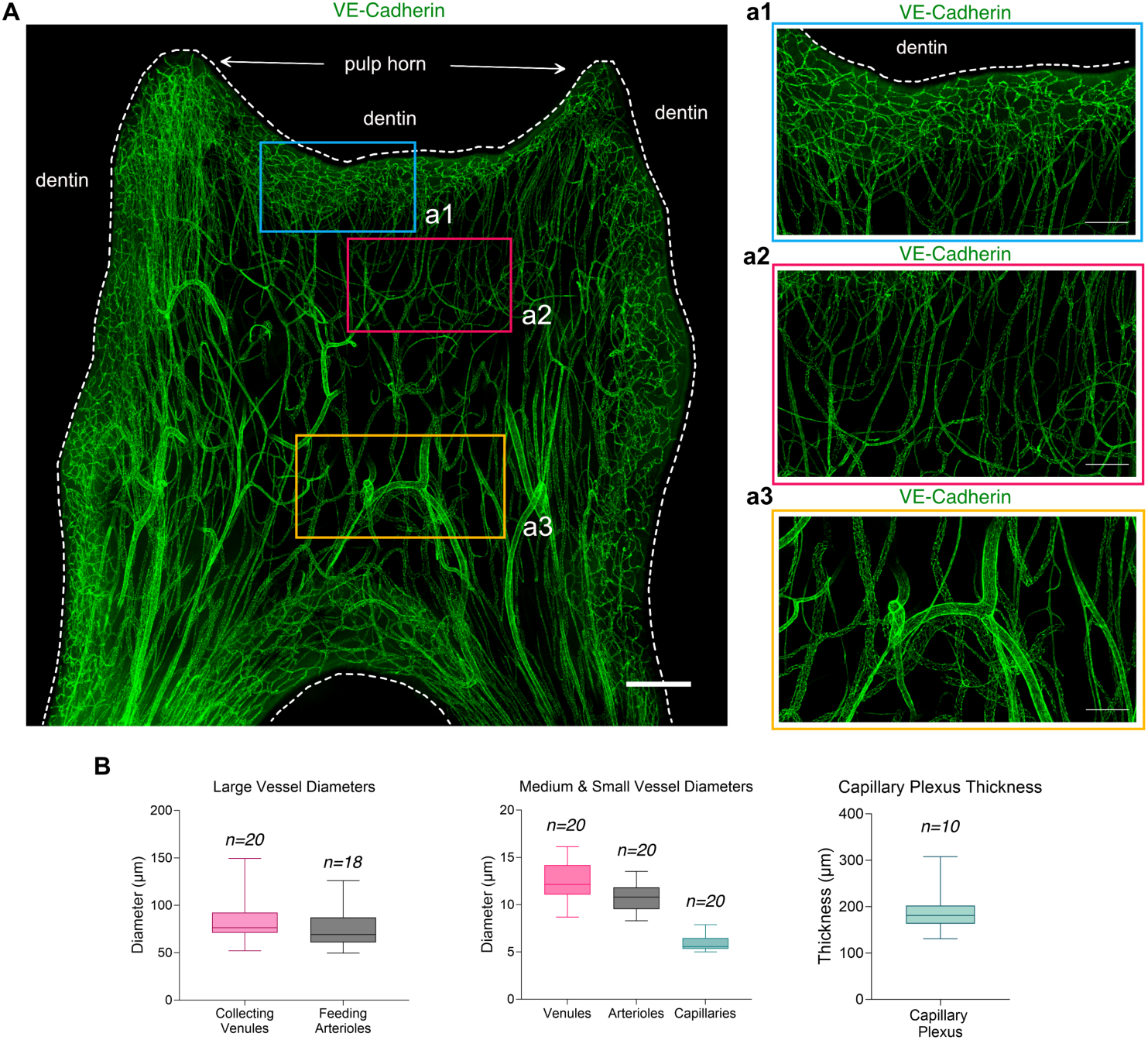
(A) Maximum intensity projection (MIP) of a 300µm z-stack showing the vasculature of a dental pulp in an immature molar showing a spatial organization in 3 layers. (a1) Zoom on the central zone containing large vessels (collecting venules and feeding arterioles) and showing a random pattern in orientation. (a2) Zoom on the intermediate zone containing medium size vessels (venules and arterioles) and showing a vertical oriented pattern. (a3) Zoom on the peripheral zone containing capillaries organized in a dense plexus, with the vessels running mostly parallel to the dentin inner surface. Nikon confocal spinning disk. Immersion objective 10x (N/A 0.5 WD5.5mm; Nikon Plan Apo). Tissue clearing using the iDISCO clearing protocol and staining for VE-Cadherin (green). Scale bar: 500µm for (A) and 200µm for (a1, a2, a3) (B) (Left) Quantification of the mean diameters of large vessels. Collecting venules and feeding arterioles have mean diameters of 84.83 μm ± 24.12 μm (n=20) and 76.08 μm ± 22.11 μm (n=18), respectively. (center) Quantification of the mean diameters of medium and small vessels. Venules, arterioles and capillaries have mean diameters of 12.46 μm ± 2.05 μm (n=19), 10.85 μm ± 1.48 μm (n=20) and 5.89 μm ± 0.74 μm (n=20); respectively. (Right) The capillary plexus has a mean thickness of 190.3 μm ± 46.64 μm (n=10).

**Figure S2.**
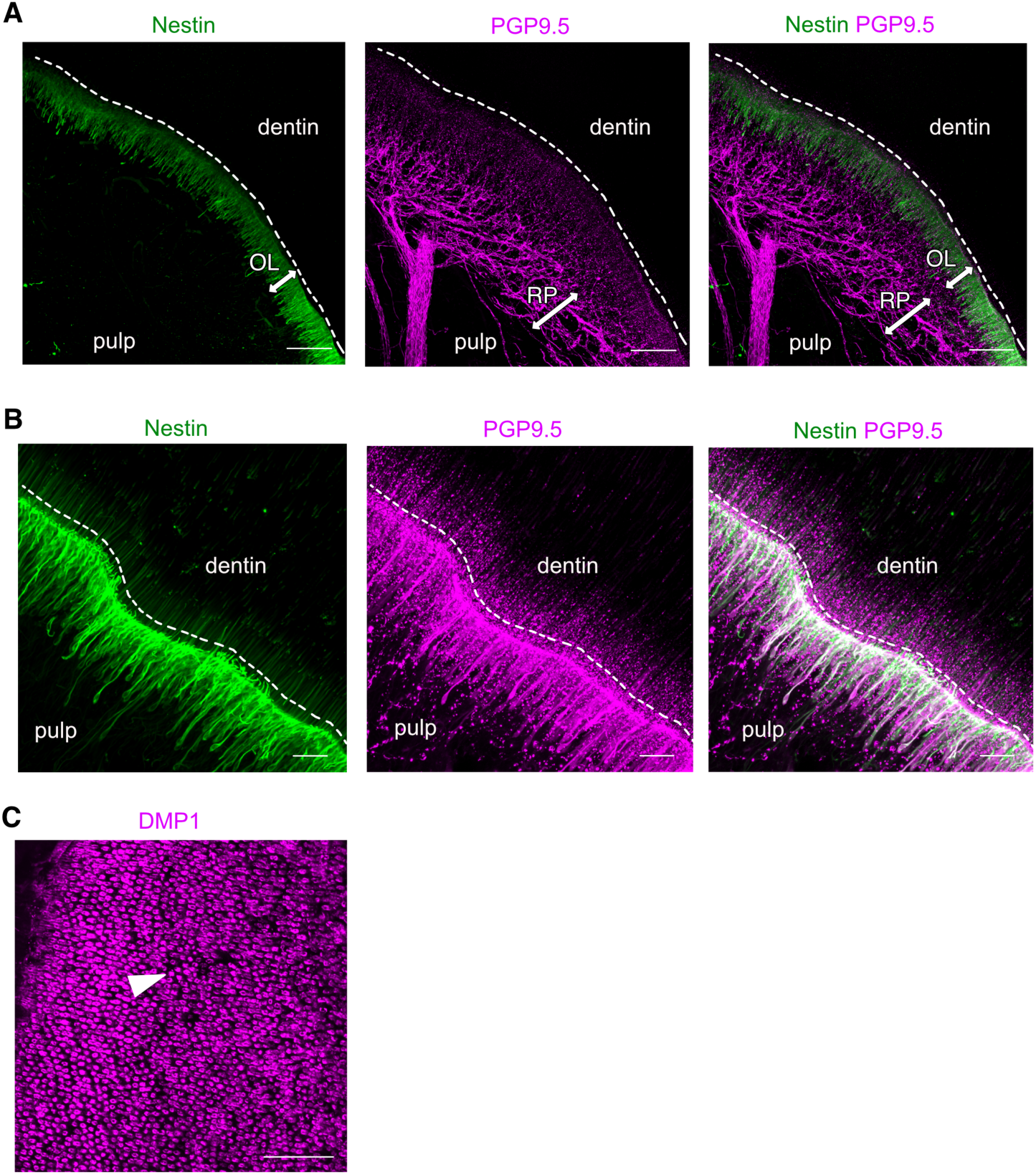
(A) Maximum intensity projection (MIP) of a 128µm z-stack showing the Raskow’s plexus and the odontoblastic layer. (B) MIP of a 91µm z-stack showing the odontoblastic layer. (C) MIP of a 18µm z-stack showing the dentinal tubules. Tissue transparization using a modified iDISCO protocol and staining for the odontoblast (A) with Nestin (green), the neuro network (B) with PGP9.5 (purple) and the dentinal tubules (C) with DMP1 (purple). Point scanning resonant confocal microscope. Plan apo Lambda 25XC Sil. optic for (A,B). Scale bars: 100µm for (A, C), 25µm for (B).

**Figure S3.**
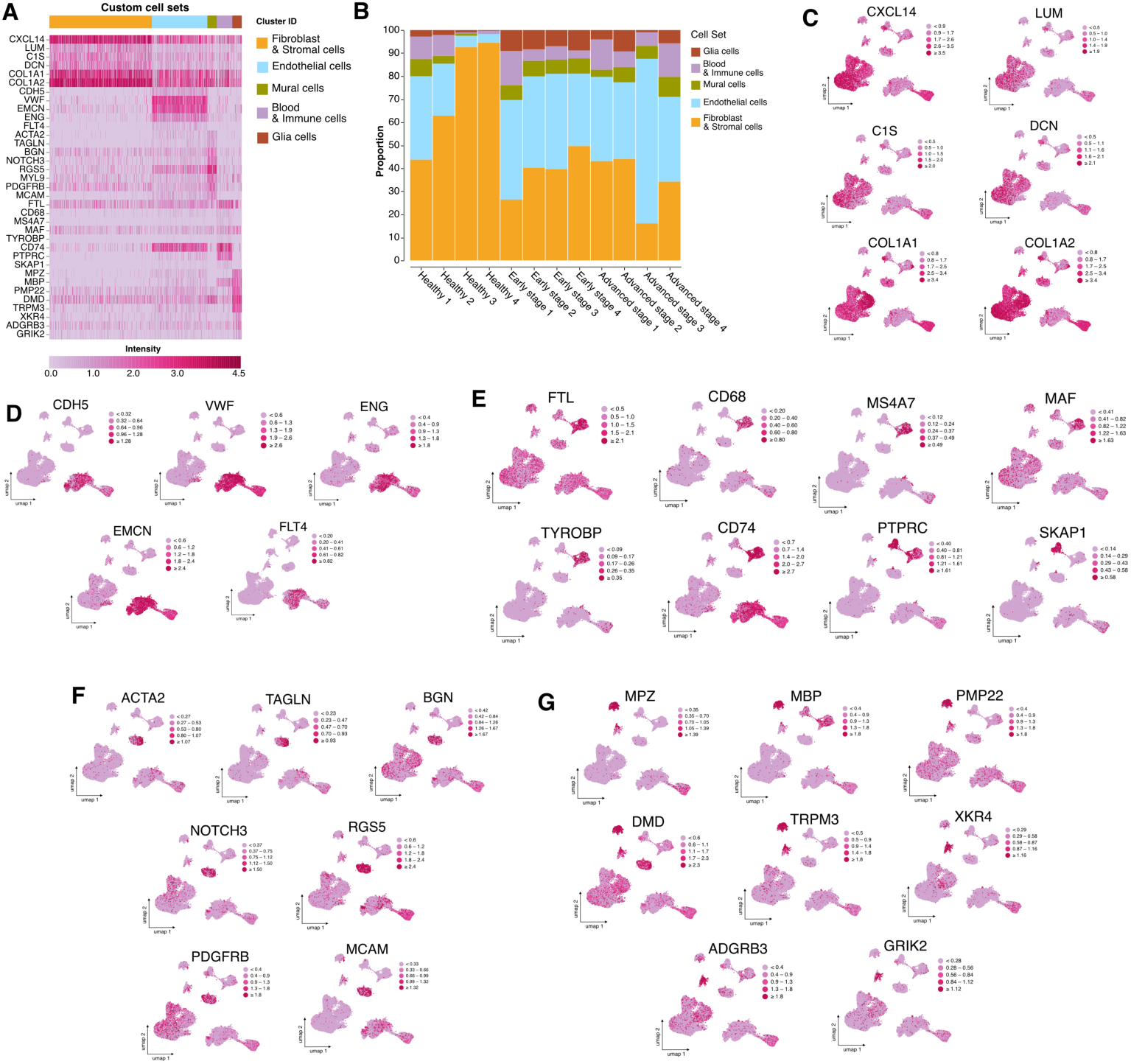
(A) Heatmap showing the relative expression of common marker gene for each cell population (B) Graph showing the proportion of each cell population across the different samples. (C) Relative expression of a well-established cell type marker for the fibroblast and stromal cells population projected on UMAP plots: *CXCL14, LUM, C1S, DCN, COL1A1, COL1A2*. (D) Relative expression of a well-established cell type marker for the endothelial cells population projected on UMAP plots: *CDH5, VWF, ENG, EMCN, FLT4*. (E) Relative expression of a well-established cell type marker for the immune cells population projected on UMAP plots: *FTL, CD68, MS4A7, MAF, TYROBP, CD74, PTPRC, SKAP1*. (F) Relative expression of a well-established cell type marker for the mural cells population projected on UMAP plots: *ACTA2, TAGLN, BGN, NOTCH3, RGS5, PDGFRB, MCAM*. (G) Relative expression of a well-established cell type marker for the glial cells population projected on UMAP plots: *MPZ, MBP, PMP22, DMD, TRPM3, XKR4, ADGRB3, GRIK2*

**Figure S4.**
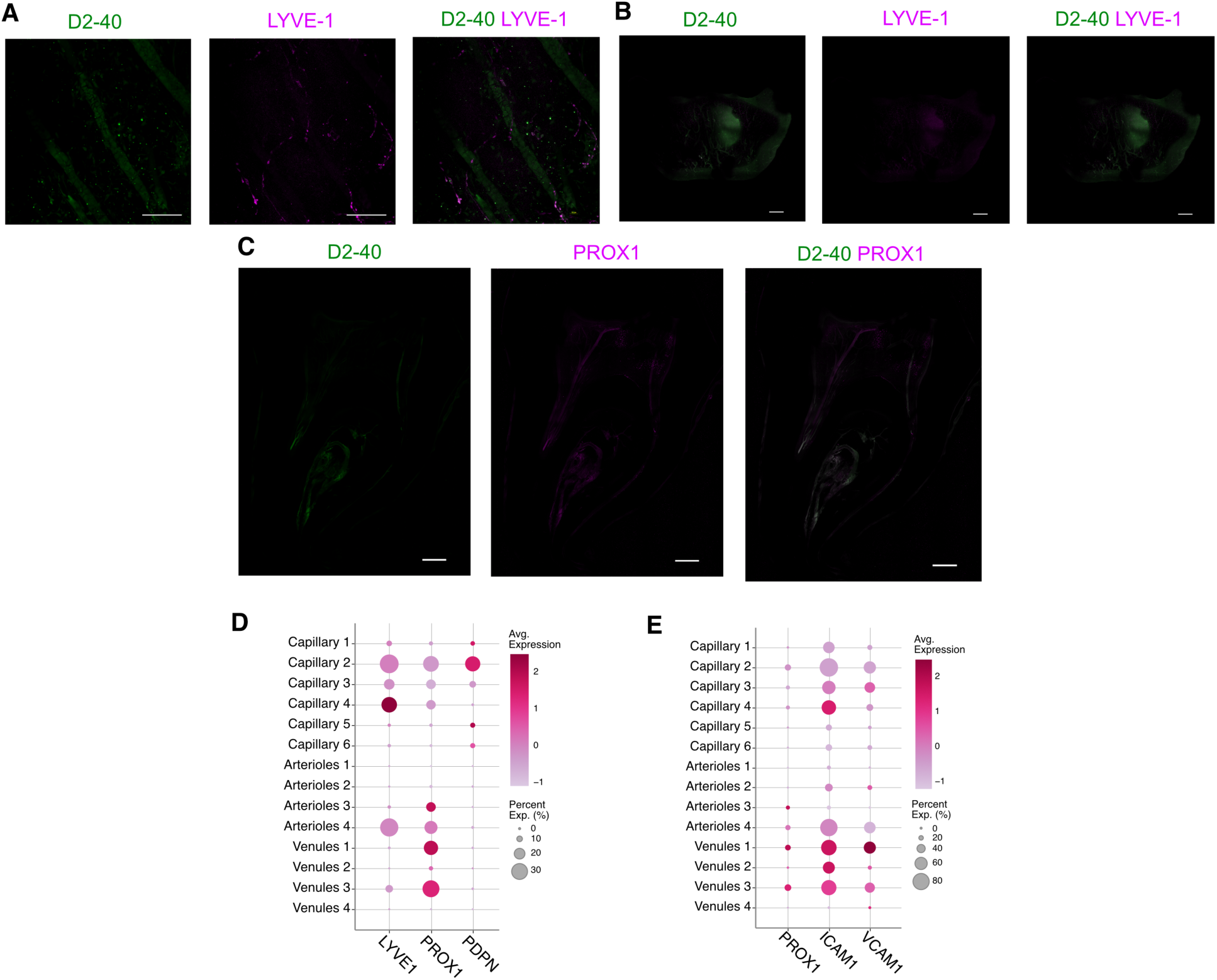
(A-C) Maximum intensity projection (MIP) of (A) a 57µm z-stack, (B) a 161µm z-stack and (C) a 141µm z-stack showing the absence of immunolabeling for lymphatic structures. Tissue transparization using a modified iDISCO protocol and staining for the lymphatic vessels (A-B) with D2-40 (green) and LYVE1 (purple) and (B) withD2-40 (green) and PROX1 (purple). Point scanning resonant confocal microscope. Plan apo Lambda 25XC Sil. optic for (A). Plan Apo 10x λS OFN25 DIC N1 optic for (B,C). Scale bars: 1000µm for (C), 500µm for (B, C), 50µm for (A). (D) Dot plot showing 3 well-established cell type markers for lymphatic vessels. (E) Dot plot showing co-expression of a well-established marker for lymphatic vessels (*PROX1*) and 2 markers associated with reactive post-capillary venules (REVs).

**Table S1.** Data on samples for single-cell experiment. This table details samples information such as name, health state, tooth number, sex, age, total cell counts and cell concentration after tissue dissociation and fixation

| Sample name | Health state | Tooth number | Sex | Age | Total cell count | Cell count/uL |
| --- | --- | --- | --- | --- | --- | --- |
| Healthy 1 | Healthy | 38 | F | 18 | 119.250 | 795 |
| Healthy 2 | Healthy | 28 | M | 18 | 228.375 | 3.045 |
| Healthy 3 | Healthy | 48 | F | 28 | 30.750 | 410 |
| Healthy 4 | Healthy | 18 | F | 28 | 24.750 | 330 |
| Early-stage 1 | Stage 1-2 | 38 | F | 19 | 83.250 | 555 |
| Early-stage 2 | Stage 2 | 38 | M | 31 | 77250 | 1.030 |
| Early-stage 3 | Stage 1-2 | 28 | F | 30 | 20.625 | 275 |
| Early-stage 4 | Stage 2 | 18 | M | 27 | 24.750 | 330 |
| Advanced stage 1 | Stage 3 | 28 | M | 29 | 28.125 | 375 |
| Advanced stage 2 | Stage 3 | 48 | M | 31 | 111.375 | 1.485 |
| Advanced stage 3 | Stage 4 | 38 | F | 30 | 25.500 | 340 |
| Advanced stage 4 | Stage 4 | 28 | M | 27 | 37.125 | 495 |

